# Sex-biased *ZRSR2* mutations in myeloid malignancies impair plasmacytoid dendritic cell activation and apoptosis

**DOI:** 10.1101/2020.10.29.360503

**Authors:** Katsuhiro Togami, Sun Sook Chung, Vikas Madan, Christopher M. Kenyon, Lucia Cabal-Hierro, Justin Taylor, Sunhee S. Kim, Gabriel K. Griffin, Mahmoud Ghandi, Jia Li, Yvonne Y. Li, Fanny Angelot-Delettre, Sabeha Biichle, Michael Seiler, Silvia Buonamici, Scott B. Lovitch, Abner Louissaint, Elizabeth A. Morgan, Fabrice Jardin, Pier Paolo Piccaluga, David M. Weinstock, Peter S. Hammerman, Henry Yang, Marina Konopleva, Naveen Pemmaraju, Francine Garnache-Ottou, Omar Abdel-Wahab, H. Phillip Koeffler, Andrew A. Lane

## Abstract

Blastic plasmacytoid dendritic cell neoplasm (BPDCN) is an aggressive leukemia of plasmacytoid dendritic cells (pDCs). BPDCN occurs at least three times more frequently in men than women, but the reasons for this sex bias are unknown. Here, studying genomics of primary BPDCN and modeling disease-associated mutations, we link acquired alterations in RNA splicing to abnormal pDC development and inflammatory response through Toll-like receptors. Loss-of-function mutations in *ZRSR2,* an X chromosome gene encoding a splicing factor, are enriched in BPDCN and nearly all mutations occur in males. *ZRSR2* mutation impairs pDC activation and apoptosis after inflammatory stimuli, associated with intron retention and inability to upregulate the transcription factor IRF7. In vivo, BPDCN-associated mutations promote pDC expansion and signatures of decreased activation. These data support a model in which male-biased mutations in hematopoietic progenitors alter pDC function and confer protection from apoptosis, which may impair immunity and predispose to leukemic transformation.

**STATEMENT OF SIGNIFICANCE:** Sex bias in cancer is well recognized but the underlying mechanisms are incompletely defined. We connect X chromosome mutations in *ZRSR2* to an extremely male-predominant leukemia. Aberrant RNA splicing induced by *ZRSR2* mutation impairs dendritic cell inflammatory signaling, interferon production, and apoptosis, revealing a sex- and lineage-related tumor suppressor pathway.

## INTRODUCTION

Blastic plasmacytoid dendritic cell neoplasm (BPDCN) is a hematologic malignancy in which patients can have leukemic involvement of the blood and bone marrow, and also tumor formation in the skin (in ~90% of cases), lymphoid organs, and in other tissues [1]. Among the unique epidemiologic features of the disease is the extreme male predominance, with a male:female incidence ratio of at least 3:1 in adults [2, 3]. There is no explanation to date for why males are predisposed to the disease, or alternatively, why females are relatively protected. Outcomes for patients with BPDCN are poor, with median survival of 12-24 months from diagnosis [3, 4], demonstrating the unmet need for additional biological insight.

DNA sequencing of bone marrow or skin involved with BPDCN has identified mutations in genes often affected in other blood cancers, particularly myelodysplastic syndrome (MDS) and related myeloid malignancies. These include point mutations or indel/frameshift mutations in *ASXL1, TET2*, and *TP53*, and copy number alterations affecting one or more cell cycle regulators [5-9]. However, BPDCN frequently arises in the context of a pre-existing or concurrent myeloid malignancy, such as MDS or chronic myelomonocytic leukemia (CMML) [10-12], but most of the prior genetic landscape studies of BPDCN analyzed unsorted bone marrow or skin biopsies with heterogeneous composition. Those biopsies likely included hematopoietic cells, possibly neoplastic, of other lineages, and non-hematopoietic cells. Therefore, one goal of this project was to define the genetic alterations and transcriptional changes present in highly purified BPDCN cells separated from any background clonal hematopoietic disorder and to link these to patient sex. We found that loss-of-function mutations in the RNA splicing factor ZRSR2, encoded on chromosome X, are enriched in BPDCN and are associated with a significant fraction of the male predominance of the disease.

BPDCN is thought to develop from plasmacytoid dendritic cells (pDCs) or their precursors, based primarily on similarities in gene expression and cellular function [13-15]. However, mechanisms of how genetic alterations promote leukemic transformation in the dendritic cell lineage are unclear. Furthermore, no studies have linked the myeloid neoplasm-associated mutations described in BPDCN to consequences on pDC development, growth, or function. Thus, a second goal of this work was to connect genes mutated in BPDCN to dendritic cell transformation via interrogation of their effects specifically in pDCs, myeloid/dendritic progenitors, and BPDCN cells. We found that *ZRSR2* mutations impair pDC activation and apoptosis in the setting of inflammatory stimuli, at least in part via misregulation of the interferon regulatory factor, IRF7, downstream of Toll-like receptor (TLR) signaling.

## RESULTS

### BPDCN has male-biased ZRSR2 mutations and a UV-associated signature

We performed whole exome sequencing (WES) on sorted BPDCN (CD45^+^ CD4^+^ CD56^+^ CD123^+^ BDCA4^+^ CD3^-^; n=11 patients) from blood or bone marrow and on paired CD3+ cells to identify acquired mutations in the malignant cells. We also performed targeted sequencing using a 95-gene panel in an extended set of bone marrows from patients with BPDCN (n=27) [16]. The most frequently mutated genes across all BPDCNs were *TET2, ASXL1*, and genes involved in RNA splicing, including *ZRSR2, SRSF2, U2AF1*, and *SF3B1* (**Figure 1a, Supplementary Table 1**). Additional mutations included genes recurrently altered in other myeloid malignancies, such as *NRAS, KRAS, TP53*, and *GNB1*. Of potential clinical relevance, we found an oncogenic *IDH2* mutation in 4 of 38 cases (~11%), which we previously reported was associated with sensitivity to the IDH2 inhibitor enasidenib in a patient with BPDCN [3]. *TET2* and *IDH2* mutations were mutually exclusive, as has been reported in acute myeloid leukemia (AML) [17]. In contrast, other myeloid malignancy-associated mutations were absent or rare, such as in *DNMT3A, NPM1*, or *FLT3-ITD* (fms-like tyrosine kinase 3, internal tandem duplication). One BPDCN harbored a single nucleotide variant that encodes an IRF8 R404W missense mutation. While the functional consequence of this specific variant is unknown, IRF8 is important for pDC development and function [18, 19]. WES also identified a small number of recurrently mutated genes in this set of BPDCNs that have not been previously identified as mutated in other cancers or in prior BPDCN sequencing (**Supplementary Table 2**). Some of these variants are in homopolymer tracts that are at risk for sequencing errors or are in genes that are not expressed in BPDCN or pDCs, and thus, their contribution to the BPDCN phenotype is not clear.

**Figure 1.**
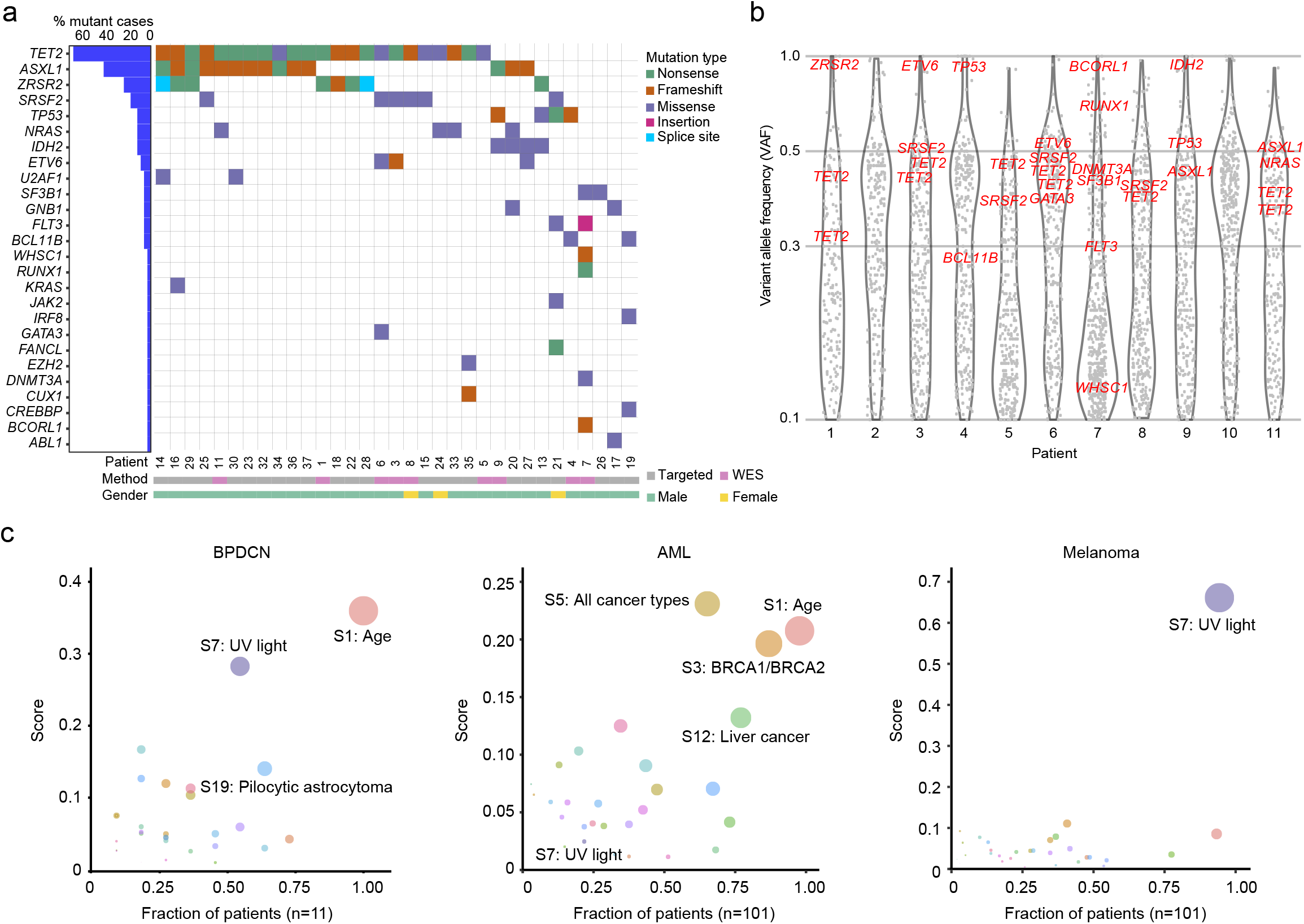
Recurrently mutated genes and ultraviolet light-induced global mutation signature in BPDCN. **a.** Co-mutation plot of single nucleotide variants (SNVs) and insertions/deletions in BPDCN samples among genes recurrently mutated in hematologic malignancies. Each column represents an individual patient, genes are in rows and mutation types are annotated by color. Percentage of patients with a given gene altered are plotted to the left. Samples are annotated by gender and sequencing method (WES, whole exome sequencing). **b.** Variant allele frequency (VAF) of all somatic mutations (including synonymous) with VAF ≥ 0.1 detected by WES in eleven BPDCNs plotted as gray dots. VAFs of known hematologic malignancy-related genes from panel (a) are annotated in red. c. Global somatic mutational signatures in BPDCN, AML, and melanoma plotted as fraction of samples (x-axis) having a specific signature with the mean signature score in those patients (y-axis). The size of the circle represents the strength of association as a combined measure of the fraction of patients having a signature and the contribution score.

In the BPDCNs studied by WES, the distribution of variant allele frequency (VAF) for all somatic mutations allowed us to define their clonal structure (**Figure 1b**). Most cases harbored known hematologic malignancy associated genes in a single dominant clone (clustered around VAF ~50% for presumed heterozygous mutations), which included all of the variants in the most frequently mutated genes: *TET2, ASXL1*, and spliceosome components. Some mutations were detected at nearly 100% VAF, including in *ZRSR2* and *BCORL1*, which reside on the X chromosome and were mutated nearly exclusively in males in this cohort, and in *ETV6, IDH2*,and *TP53*, which likely represent loss-of-heterozygosity events.

Given that many genes mutated in BPDCN were shared with other myeloid malignancies, we asked if the global mutation pattern might indicate unique BPDCN-specific features. We performed a mutation analysis on all somatic alterations and their surrounding nucleotide context using signatures previously defined in the Catalog of Somatic Mutations in Cancer (COSMIC) [20]. We found that BPDCNs harbored an age-associated pattern (“Signature 1”), which is detected in most cancers in COSMIC (**Figure 1c, Supplementary Figure 1**). However, we also detected a strong association with ultraviolet (UV)-induced mutation signatures in the majority of BPDCNs (signature score >0.25 in 6/11 or 55%; **Figure 1c**). In contrast, there was no UV signature associated with AML, even though AML shares many recurrently mutated genes with BPDCN and may arise from a similar myeloid progenitor. For comparison, melanoma harbors a uniformly high UV signature in most tumors (**Figure 1c**). The BPDCN cells we analyzed were harvested from bone marrow or blood and not directly from UV-exposed tissue (i.e., skin), suggesting that at least some leukemic BPDCN cells maintain a UV signature presumably acquired during a prior skin phase of malignant evolution.

Sequencing purified BPDCN cells rather than bulk marrow or skin samples also allowed us to determine DNA copy number variants (CNVs) that were more confidently BPDCN-associated. Some of the chromosome and arm level CNVs were similar to those described previously in BPDCN (**Figure 2a**) [7, 21]. These included loss of 7p (which harbors *IKZF1)*, 9p *(CDKN2A, CDKN2B)*, 12p *(CDKN1B, ETV6)*, 13q (*RB1*), and 17p (*TP53*) (**Supplementary Figure 2**). Copy number analysis also clarified the disease genetics for patients 2 and 10, who did not have any single nucleotide variants in known blood cancer-associated genes from WES. In those patients, we detected deletions in *IKZF1* and *RB1* (patient 2) and *TET2, RB1*, and *ZRSR2* (patient 10), which supports that analysis of CNVs in addition to point mutations/indels can detect additional relevant driver events in patients. Integrated analysis of single nucleotide variants/indels and copy loss highlighted four putative BPDCN tumor suppressors targeted by both types of alteration: *ZRSR2, TET2, TP53*, and *SETD2* (**Figure 2b**). PCR confirmed the WES finding that chromosome Xp22.2 copy loss in patient 10, a male, resulted in complete absence of *ZRSR2* DNA in tumor but not in germline cells (**Figure 2c**).

**Figure 2.**
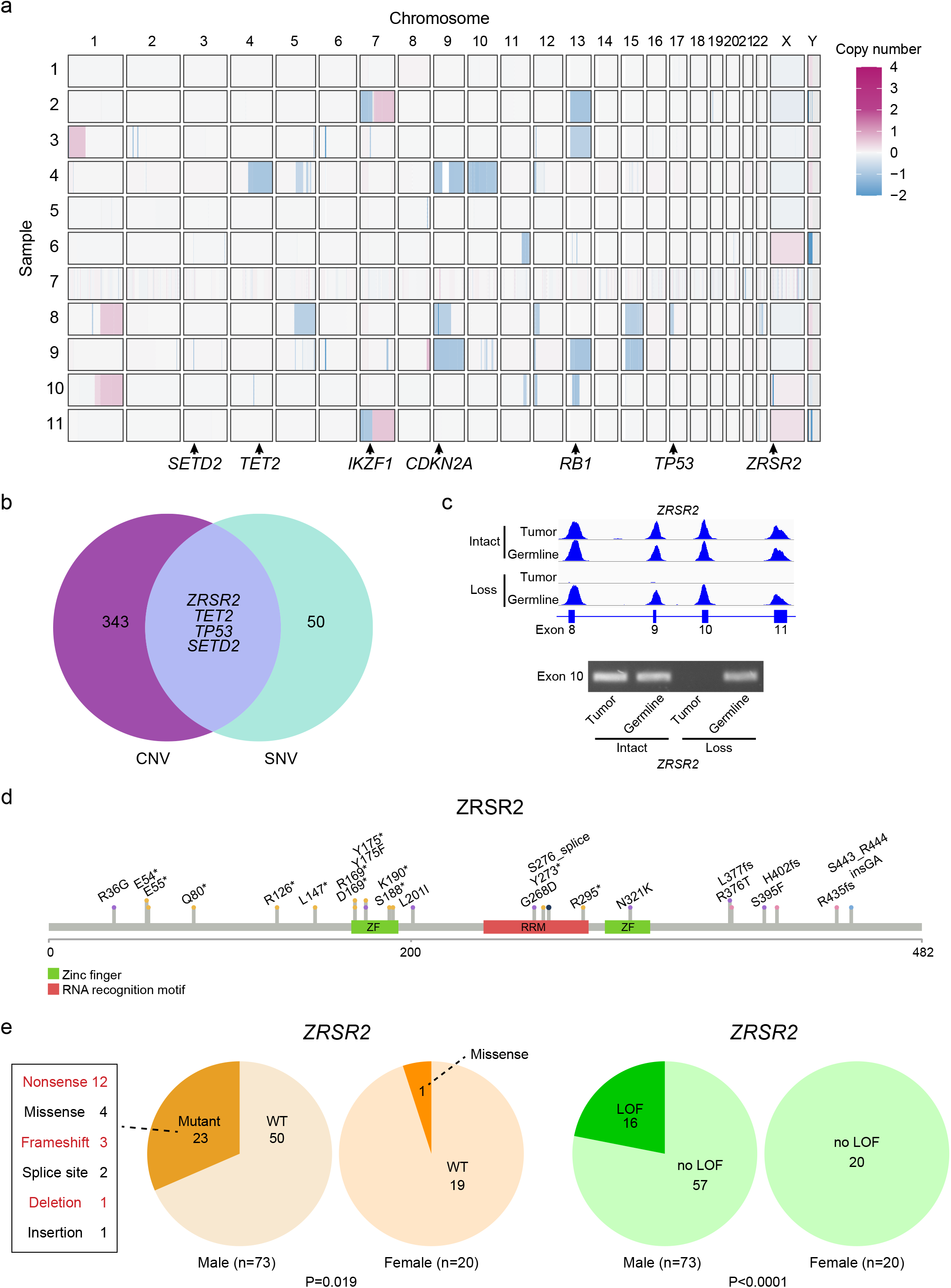
*ZRSR2* mutations and their association with BPDCN sex bias. **a.** Somatic DNA copy number changes in 11 BPDCNs identified by WES shown from blue (copy loss) to red (gain). **b.** Venn diagram showing the four genes with overlap of copy number variations (among 347 protein coding genes observed in at least two patients with VAF ≥ 0.2) and single nucleotide variations (among 54 targeted panel genes or protein coding gene mutations observed in at least two patients by WES). **c.** Sequencing read traces in the *ZRSR2* locus from WES and PCR of exon 10 DNA in tumor/germline pairs from representative BPDCNs in males with intact or somatic copy number loss of *ZRSR2*. **d.** Schematic of the ZRSR2 protein with amino acid locations and specific mutations (n=23) detected in BPDCN (n=93). **e.** *ZRSR2* mutations are male-biased. 23 of 73 male BPDCNs had *ZRSR2* mutations vs. 1 of 20 female (P=0.019 by Fisher’s exact test). When restricted to obvious loss-of-function (LOF) mutations (nonsense, frame shift, deletion; marked in red), 16 of 73 male BPDCNs had *ZRSR2* LOF mutations vs. 0 of 20 female (P<0.0001).

*ZRSR2* mutations in BPDCN are of particular interest for several reasons. First, while other splicing factors are mutated across many solid and blood cancers, *ZRSR2* has been associated with myeloid disorders, such as MDS, CMML, and AML. However, in contrast to MDS, where *SRSF2, SF3B1*, and *U2AF1* mutations are more common, *ZRSR2* was the most frequently mutated spliceosome gene in this BPDCN cohort. To confirm this finding in a larger and more geographically diverse cohort, we examined BPDCN DNA from three separate groups (from France, Italy, and MD Anderson Cancer Center), and found *ZRSR2* mutations in 24 of 93 (26%) (**Figure 2d**). This is significantly higher than the incidence of *ZRSR2* mutations in MDS, where the gene was mutated in only 13 of 288 patients in one large cohort (4.5%; P<0.0001 by Fisher’s exact test) [22].

Second, nearly all *ZRSR2* mutations in our cohort were in males (mutated in 23 of 73 males vs. 1 of 20 females, P=0.019 by Fisher’s exact test) (**Figure 2e**). The mutational pattern is consistent with loss-of-function, with most variants predicting inactivating events (nonsense, frame shift, deletion), which implicates *ZRSR2* as a male-biased X chromosome tumor suppressor gene. In fact, strictly defined inactivating mutations in our cohort were exclusively in males (16 of 73 males vs. 0 of 20 females, P<0.0001 by Fisher’s exact test) (**Figure 2e**). This is similar to MDS, where loss-of-function mutations in *ZRSR2* are almost always restricted to males [22]. *ZRSR2* belongs to the minor subset of X chromosome genes that escape X inactivation silencing and is biallelically expressed in female cells [23]. The finding that *ZRSR2* is also preferentially mutated in PBDCN and MDS from males strongly suggests that *ZRSR2* is an Escape from X Inactivation Tumor Suppressor (EXITS) gene [24]. As we previously described for other EXITS genes across a range of cancers, females are relatively protected from BPDCN because they have two active *ZRSR2* alleles and therefore require two mutations to eliminate function, whereas males require only one. The degree of sex bias in BPDCN incidence linked to *ZRSR2* can be expressed as the number of excess *ZRSR2* mutations in males per excess case of BPDCN in males [24]. Using that calculation, in this cohort ~42% of the excess male risk of BPDCN is associated with a *ZRSR2* mutation.

### Unique transcriptomic features of spliceosome mutated BPDCN

Next, we asked how BPDCN transcriptomes relate to their genetics. We performed RNA-sequencing on the same sorted BPDCN samples that we had profiled by whole exome sequencing. Peripheral blood pDCs (CD45^+^ CD123^+^ BDCA2^+^ CD3^-^) from healthy donors were analyzed in parallel for comparison (**Supplementary Table 3**). Differentially expressed genes between BPDCN and pDCs were similar to those we previously defined that distinguished BPDCNs from normal dendritic cells using single cell RNA-sequencing (**Figure 3a**) [14]. We also observed differences in individual oncogenes and tumor suppressors (*BCL2, MYB, IRF4*, and *MAP3K1*) known to be dysregulated in BPDCN (**Figure 3b**). Furthermore, gene set enrichment analysis (GSEA) identified oncogene-associated signatures in BPDCN, including overexpression of MYC and E2F targets, and of PI3K/AKT/MTORC1 signaling pathway genes (**Figure 3c**). In addition to these expected gene expression changes, we also found that BPDCNs had upregulation of RNA splicing machinery and associated genes when compared to normal pDCs (**Figure 3d**). Furthermore, within BPDCN, splicing factor mutant cases showed upregulation of RNA splicing genes and also were enriched for markers of active nonsense mediated decay (NMD) RNA catabolism compared to BPDCNs without *ZRSR2, SRSF2, SF3B1*, or *U2AF1* mutations (**Figure 3d**). This was of particular interest given that ZRSR2 loss promotes intron retention [25], which is known to trigger NMD activation and cause altered myelopoiesis [26].

**Figure 3.**
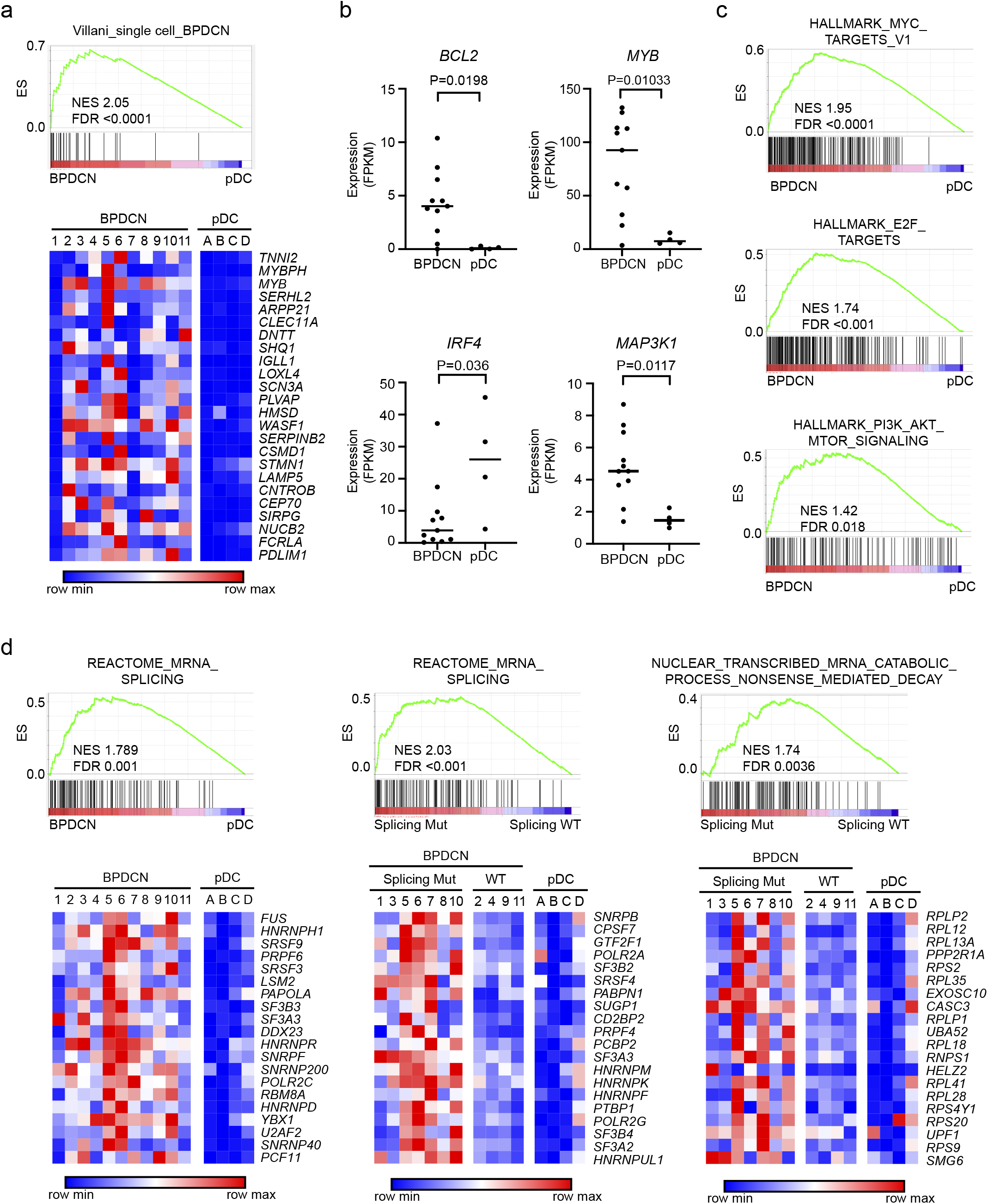
BPDCN transcriptomes have alterations in oncogene, dendritic cell development, and RNA processing genes. **a.** Gene set enrichment analysis (GSEA) showing association of a previously-defined gene signature from single cell RNA-sequencing that differentiated BPDCN from normal human DC subtypes [14] in BPDCN (n=11) compared to normal pDCs (n=4) from the current cohort. ES, enrichment score; NES, normalized enrichment score; FDR, false discovery rate. Heatmaps of the same genes plotted as low (blue) to high (red) relative expression. **b.** RNA expression of the indicated genes in BPDCN (n=11) compared to normal pDCs (n=4), groups compared by t test. **c.** GSEA comparing BPDCN to normal pDCs showing enrichment of the indicated hallmarks of cancer gene sets (MSigDB collection “H”) in BPDCN. **d.** GSEA as in (a) for the indicated RNA splicing and nonsense mediated decay gene sets with heatmaps of the top 20 leading edge genes plotted as low (blue) to red (high) relative expression. In the indicated plots, BPDCNs are separated by whether (Splicing Mut) or not (Splicing WT) they harbor a mutation in a splicing factor (*SF3B1, SRSF2, U2AF1*, or *ZRSR2*).

Next, we analyzed RNA splicing in splicing factor mutated BPDCN compared to normal pDCs and to BPDCN without any splicing factor mutation. BPDCNs with splicing factor mutations showed several types of abnormal splicing, including intron retention (IR), cryptic 3’ splice site usage, and exon skipping (**Supplementary Figure 3a, Supplementary Table 4**). ZRSR2, SRSF2, SF3B1, and U2AF1 are all involved in U2 intron splicing (99.7% of introns in the human genome). In contrast, ZRSR2 is uniquely necessary for proper splicing of U12 introns (0.3% of human introns), an evolutionarily conserved genomic feature with distinct branch and splice-site sequences [27, 28]. In MDS, *ZRSR2* mutations promote aberrant IR with a bias toward retention of U12-type introns [29]. Similarly, we found that BPDCNs with *ZRSR2* mutation or loss had increased IR compared to those without spliceosome mutations (**Figure 4a**). Both U2- and U12-type introns were affected, but IR was markedly weighted towards U12-type introns in BPDCNs with *ZRSR2* mutation (**Figure 4b**). We also detected some of the stereotypical mis-splicing patterns associated with specific splicing factor mutations in other cancers [29]. These included preferential exon inclusion or exclusion related to CCNG/GGNG exonic splicing enhancer motifs in cases with *SRSF2* mutation [30] and alternative 3’splice sites usage with *SF3B1* mutation (**Supplementary Figure 3a-b**) [31]. To confirm ZRSR2-associated splicing changes in a separate set of BPDCNs, we analyzed patient-derived xenografts (PDXs) [32]. We performed RNA-seq on six BPDCN PDXs, two with *ZRSR2* mutations and four without. Similar to what we observed in patient cells, PDXs with *ZRSR2* mutations had increased IR events that were biased toward aberrant retention of U12-type introns (**Figure 4b-c**).

**Figure 4.**
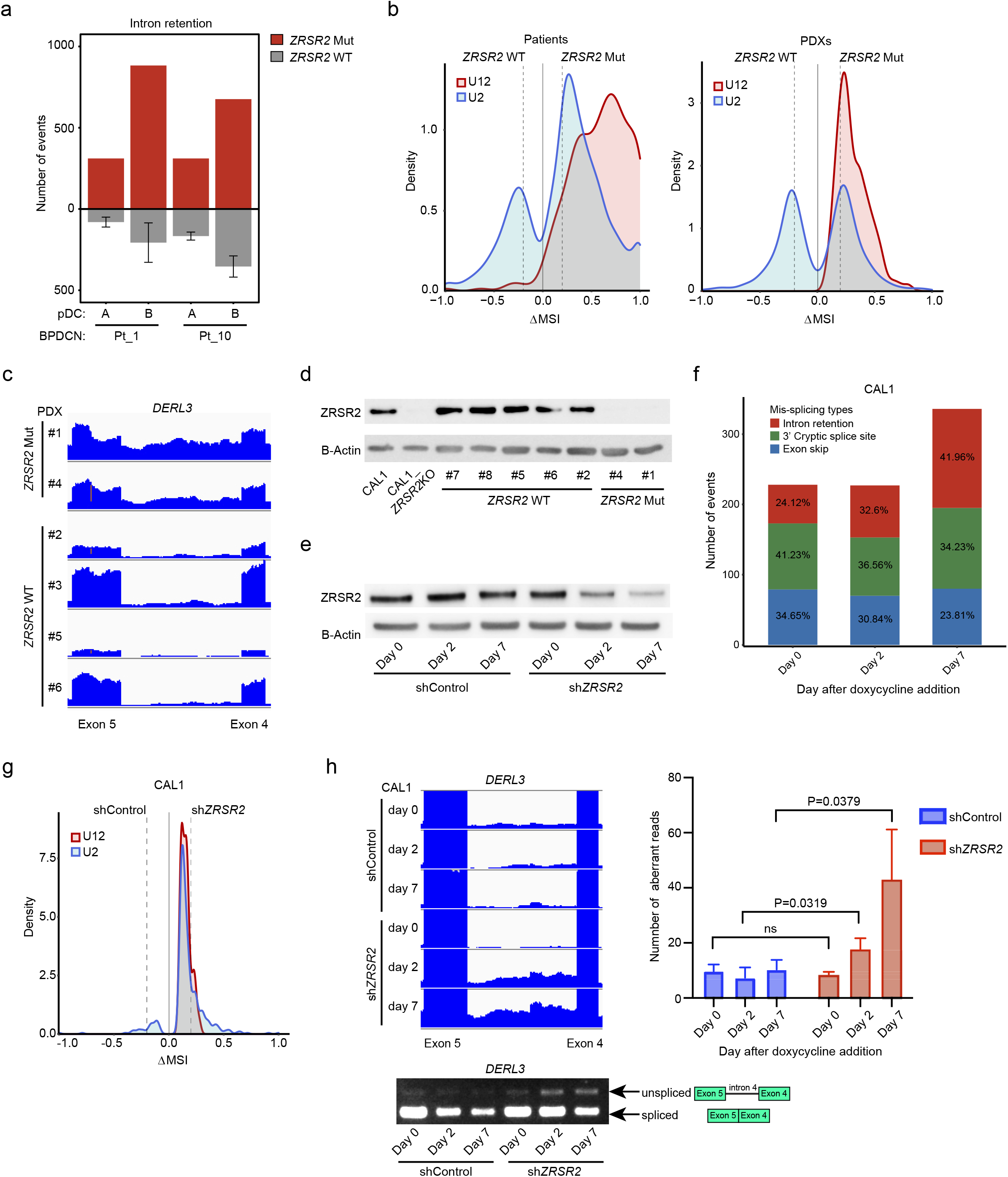
ZRSR2 mutations promote specific mis-splicing events. **a.** Intron retention events in BPDCNs with a *ZRSR2* mutation (n=2, red) compared pairwise to normal pDCs (n=2, noted as samples A and B) and to those events in BPDCNs with no known splicing factor mutation (n=4, gray). **b.** Density plots of frequencies of U2-type (blue) and U12-type (red) introns among aberrantly retained introns (Fisher’s exact test p-value ≤ 0.05) in pairwise analyses of *ZRSR2* mutant versus wild-type BPDCNs and patient-derived xenografts (PDXs). Dotted lines represent ΔMSI −0.2 and +0.2. **c.** RNA-seq reads of an aberrantly retained U12 intron (intron 4) in human *DERL3* from BPDCN PDXs with or without *ZRSR2* mutation (equivalent scale for all samples). **d.** Western blot for ZRSR2 and beta-actin in CAL1 cells (parental or ZRSR2 knockout) and in BPDCN PDXs with wild-type or mutant *ZRSR2*. **e.** Western blot from CAL1 cells at baseline (day 0) and at 2 and 7 days after doxycycline induction of non-targeting control or *ZRSR2*-targeted shRNAs. **f.** Mis-splicing events after knockdown of *ZRSR2* from cells shown in panel (e), calculated by pairwise comparison between ZRSR2 knockdown and controls (n=3 biological replicates each) on days 0, 2, and 7 of doxycycline induction. The number of events on each day is depicted in a single bar colored by different types of mis-splicing events: intron retention (red), 3’ cryptic splice site (green) and exon skip (blue). **g.** Density plots of frequencies of U2-type (blue) and U12-type (red) introns in pairwise analysis of CAL1 with *ZRSR2* knockdown versus control on day 7 after the doxycycline induction (Fisher’s exact test p-value ≤ 0.05), plotted as in (b). **h.** (left) RNA-seq reads and RT-PCR from CAL1 cells after the induction of control or *ZRSR2*-targeted shRNA showing retention of *DERL3* intron 4. (right) Number of *DERL3* intron 4 RNA-seq reads plotted for control and *ZRSR2* knockdown cells at days 0, 2, and 7 after doxycycline shRNA induction (n=3 biological replicates each, groups compared by t test).

Next, we asked if the splicing abnormalities observed were directly related to loss of *ZRSR2*. The predominant nonsense and frameshift mutations we detected in BPDCN (**Figure 2d-e**) are predicted to promote nonsense mediated RNA decay and absence of a mature protein, which is what we observed by western blotting in BPDCN with *ZRSR2* mutations (**Figure 4d**). Therefore, we used engineered knockdown or knockout cell line models to study *ZRSR2* mutations in BPDCN cells. First, we generated BPDCN cell lines harboring doxycycline-inducible shRNAs targeting *ZRSR2* to model protein loss and facilitate time-dependent analysis of ZRSR2 depletion effects on splicing (**Figure 4e**). Seven days after induction of knockdown, intron retention was the most prominent aberrant splicing event in *shZRSR2* compared to shControl cells (**Figure 4f**). Both U2- and U12-type intron retention events were increased in ZRSR2 knockdown cells (**Figure 4g**) and intron retention events detected by RNA-seq could be confirmed by RT-PCR (**Figure 4h**). These experiments suggested that U2- and U12-type intron retention is a direct and immediate consequence of ZRSR2 depletion on RNA splicing in human BPDCN cells.

### ZRSR2 mutation impairs pDC activation and apoptosis induced by inflammatory stimuli

Genes that were misspliced in primary BPDCNs with *ZRSR2, SRSF2*, or *SF3B1* mutations included several that are important in dendritic cell development and/or function, including *CSF2RB, IRF4, IRF7, IRF8*, and *LILRB4* (**Supplementary Table 4**). Therefore, to connect missplicing with cellular phenotype we asked how *ZRSR2* mutations affect pDC function. *ZRSR2* knockdown was inversely correlated with expression of genes normally upregulated in Toll-like receptor 7 (TLR7) agonist (R848, resiquimod) treated dendritic cells (**Figure 5a**). This was of particular interest because studies have reported that primary BPDCN cells are less responsive to TLR/infectious stimulation than normal pDCs or have signatures of decreased activation, but the mechanisms are unknown [13, 33]. To test if complete loss of ZRSR2, as we saw in patient samples, was sufficient to impair pDC activation, we generated BPDCN CAL1 cells with CRISPR/Cas9 knockout of *ZRSR2* (**Figure 5b**). We then stimulated the knockout cells with TLR agonists. Upregulation of the activation marker CD80 after stimulation with lipopolysaccharide (LPS, TLR4 agonist) or R848 (TLR7 agonist) was markedly reduced in lines with knockout of *ZRSR2* (**Figure 5c**). TLR stimulation of normal pDCs induces secretion of numerous inflammatory cytokines, including type 1 interferons (IFN-a and IFN-β). In contrast, CAL1 cells with *ZRSR2* knockout had defective secretion of several cytokines (e.g., IFN-a, IFN-β, IL-6, TNF-a) after LPS or R848 stimulation compared to controls (**Figure 5d**). We also found similar defective cytokine production in primary BPDCN cells compared to normal pDCs upon stimulation with R848 (**Figure 5d**). Of note, ZRSR2 loss did not simply cause a complete block in TLR signaling or protein secretion, because other cytokines were produced equivalently in ZRSR2-mutant cells and BPDCN cultures after stimulation (e.g., TRAILR1). Together, these data suggest that *ZRSR2* mutations in BPDCN cells cause defective activation and impaired secretion of specific cytokines, including type 1 interferons, in the setting of TLR stimulation.

**Figure 5.**
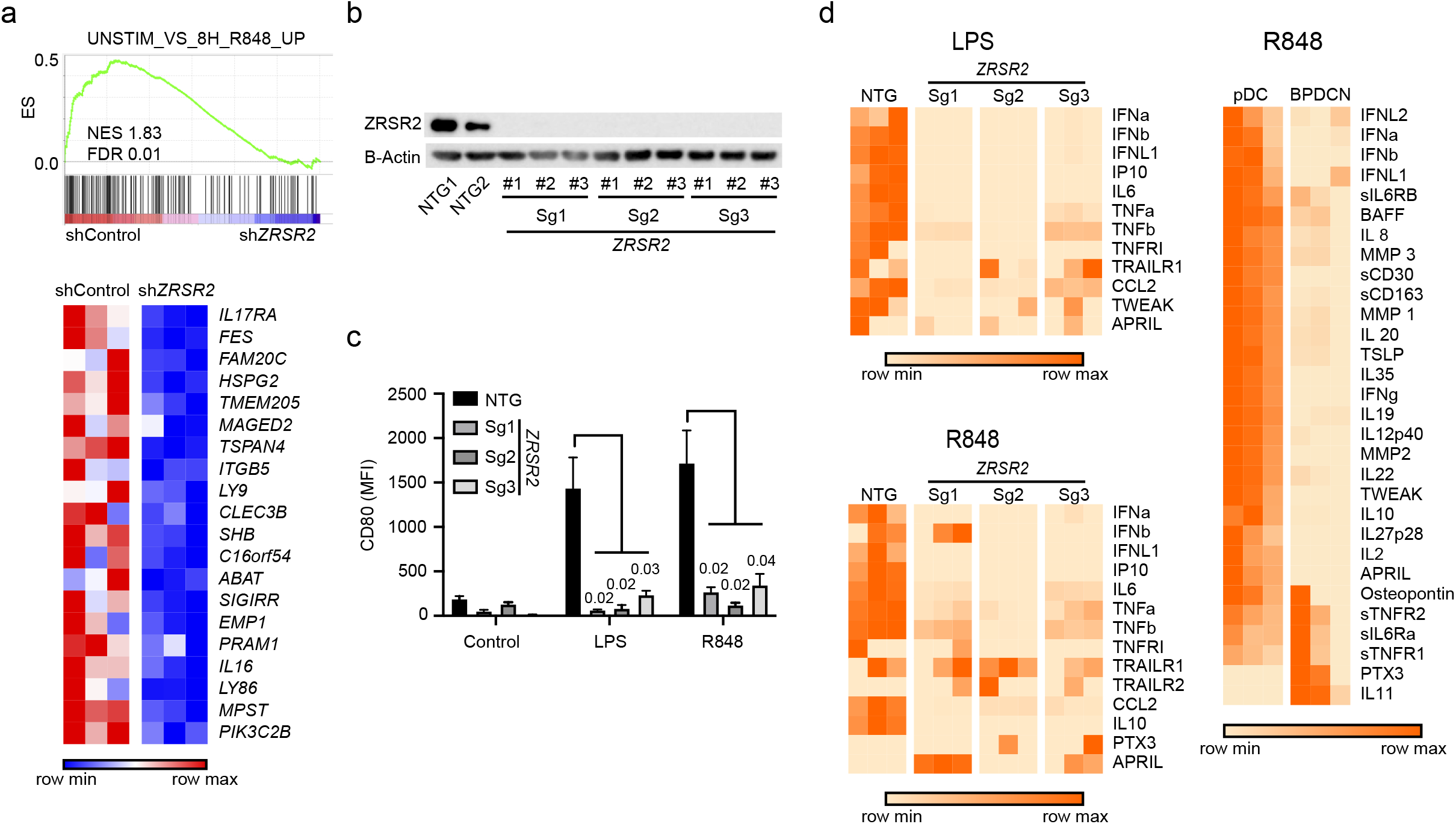
pDC activation by Toll-like receptor stimulation is impaired by loss of ZRSR2. **a.** GSEA of RNA-seq showing decreased enrichment of TLR7 (R848) stimulated genes in *ZRSR2* knockdown cells. Heatmap shows expression levels of the leading edge genes in GSEA from blue (low) to red (high). ES, enrichment score; NES, normalized enrichment score; FDR, false discovery rate. **b.** Western blot for ZRSR2 and beta-actin in control or ZRSR2 knockout CAL1 cells. NTG1 and NTG2 are independent non-targeting sgRNAs and Sg1, Sg2, and Sg3 are independent *ZRSR2*-targeted sgRNAs, each assessed in biological triplicate samples. **c.** Mean fluorescence intensity of cell surface CD80 on control or ZRSR2 knockout CAL1 cells is shown after stimulation with either LPS or R848 (n=3 biological replicates of each sgRNA, groups compared by t test). **d.** Heatmaps showing protein quantitation of the indicated cytokines in supernatants of control or ZRSR2 knockout CAL1 cells (n=3 independent biological replicates of each sgRNA, non-targeting or *ZRSR2*-targeting), and normal pDCs or BPDCN PDXs (each column is from an independent individual donor or PDX) after stimulation with LPS or R848.

After activation, normal pDCs undergo apoptotic cell death as part of a negative feedback process that limits inflammation [34]. We hypothesized that *ZRSR2* mutations might protect pDCs from apoptosis in the setting of TLR stimulation and this could be a mechanistic link between spliceosome alterations and malignant transformation to BPDCN. We stimulated CAL1 cells with LPS or R848 and found that both initiated apoptosis as measured by activation of caspase 3 and 7 (**Figure 6a**). Knockout of *ZRSR2* conferred relative protection from LPS or R848-induced apoptosis (**Figure 6b**). Interestingly, the growth rate of CAL1 cells at steady state was not changed by *ZRSR2* mutation. In contrast, while growth of wild-type cells was inhibited by LPS, *ZRSR2* knockout cells were protected from LPS-induced growth arrest (**Figure 6c**).

**Figure 6.**
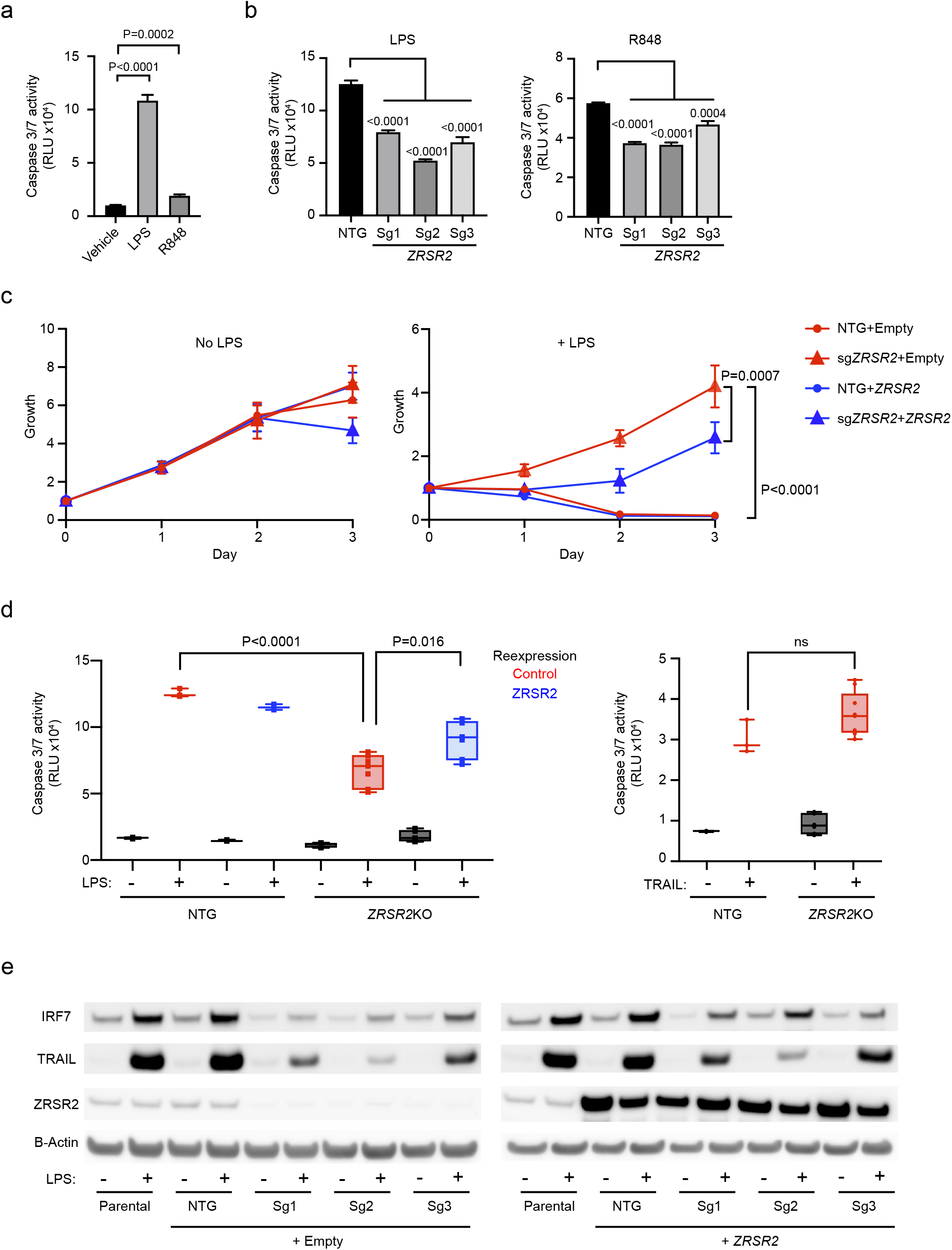
ZRSR2 mutation impairs pDC apoptosis following TLR stimulation associated with blunted IRF7 and TRAIL induction. **a.** Caspase 3/7 activity in parental CAL1 cells after treatment with LPS or R848 compared to vehicle control. **b.** Caspase 3/7 activity in control or ZRSR2 knockout cells after stimulation with LPS or R848 compared to control sgRNA-expressing cells. **c.** Relative growth of control and ZRSR2 knockout cells, with wild-type ZRSR2 reexpression or empty vector control, are shown in normal medium (left) or in medium containing LPS (right). **d.** Caspase 3/7 activity in control and ZRSR2 knockout cells, with wild-type ZRSR2 reexpression or empty vector control, is shown after treatment with LPS or TRAIL. In panels a-d, n=3 biologically independent replicates, groups compared by t test. **e.** Western blot for IRF7, TRAIL, ZRSR2 and beta-actin in parental, nontargeting control, and ZRSR2 knockout CAL1 cells, with or without ZRSR2 reexpression, 24 hours after stimulation with LPS or vehicle.

We confirmed the specificity of CRISPR knockout and that loss of ZRSR2 was necessary for protection from apoptosis by demonstrating that reexpression of wild-type ZRSR2 in mutant cells partially rescued the ability of LPS to induce apoptosis and impair growth (**Figure 6c-d**). Finally, we confirmed that *ZRSR2*-mutant CAL1 cells had not simply lost the ability to undergo TLR activation-induced cell death. A downstream consequence of pDC stimulation is production of TRAIL, a TNF-family pro-apoptotic cytokine, which promotes autocrine and paracrine apoptosis [35, 36]. Treatment with exogenous TRAIL promoted apoptosis equivalently in control and *ZRSR2* mutant cells, demonstrating that the cell death response downstream of TRAIL was intact (**Figure 6d**).

Collectively, these data suggested that *ZRSR2* mutation may perturb specific pathways downstream of TLRs that result in impaired activation-induced cell death. While it is likely that multiple effects of splicing mutations contribute to BPDCN, we asked if any common targets across BPDCNs might be involved in the hypo-activation phenotype. When we analyzed aberrant splicing across BPDCNs harboring any splicing factor mutation, *IRF7* was among the overlapping candidates. The *IRF7* gene encodes a transcription factor, interferon regulatory factor 7, that is activated by TLR signaling and is important for induction of downstream genes, including type 1 interferons and TRAIL [37, 38]. Interestingly, the *IRF7* mRNA transcript is not classified as having U12-type introns but it does contain a so-called “weak intron” (intron 4) that is known to be subject to intron retention and NMD in normal dendritic cells during activation [39]. Rate-limiting splicing is a feature of U12-type introns and also more generally of a group of genes (with either U2 or U12 introns) that participate in “programmed delayed splicing” associated with inflammatory response regulation at the level of RNA processing [40, 41]. *IRF7* intron 4 was aberrantly retained in *ZRSR2* mutant BPDCNs (2 of 2) and the same intron-exon region was also misspliced in *SRSF2* (4 of 4) and *SF3B1*-mutated (1 of 1) cases but not in splicing factor wild-type BPDCNs or in normal pDCs (**Supplementary Table 4**).

Therefore, we hypothesized that TLR stimulation of BPDCN with a *ZRSR2* mutation would result in impairment of IRF7 induction and downstream gene activation including pro-apoptotic pathways. Indeed, whereas the baseline IRF7 protein levels were minimally affected by *ZRSR2* mutation, mutant cells had severely impaired ability to increase IRF7 protein after LPS stimulation (**Figure 6e**). *IRF7* mRNA increased, as expected, after stimulation in both control and mutant cells (**Supplementary Figure 4a**), which supports that the impairment of IRF7 protein induction by LPS is post-transcriptional. Induction of TRAIL was also impaired in *ZRSR2*-mutant cells, as expected for an IRF7 target gene. IRF7 and TRAIL induction by LPS were partially rescued in *ZRSR2*-mutant cells by overexpression of wild-type ZRSR2 (**Figure 6e**). Furthermore, expression of an intronless, constitutively active IRF7 [42] impaired growth of both wild-type and *ZRSR2*-mutant cells (**Supplementary Figure 4b**). This suggested that loss of ZRSR2 affected the induction of IRF7 protein after LPS stimulation, but activity downstream of activated IRF7 remained intact. Together, these data support a model in which *ZRSR2* mutation in BPDCN inhibits apoptosis and promotes a growth advantage compared to non-mutated cells in the setting of TLR activation.

### BPDCN-associated Zrsr2 and Tet2 mutations promote aberrant pDC phenotypes in vivo

To evaluate the contribution of BPDCN-associated mutations to pDC development and function in vivo, we generated bone marrow chimeras with hematopoietic cells harboring mutations in *Zrsr2, Tet2*, or both. *Tet2* was chosen because it is the most frequently mutated gene in BPDCN and is the most commonly co-occurring mutated gene in BPDCNs with mutated *ZRSR2*. We harvested c-kit^+^ bone marrow cells from Rosa-Cas9 knock-in animals, transduced with lentiviruses encoding sgRNAs targeting *Zrsr2, Tet2*, or control non-targeting guides in pairwise fashion, each co-expressing GFP or tagRFP. Eight weeks after transplantation of transduced marrow into lethally irradiated wild-type recipients, we observed approximately equivalent single- and double-positive GFP/tagRFP cells in the bone marrow across genotypes (**Figure 7a**). We confirmed CRISPR insertion-deletion events at sgRNA target sites using a T7 endonuclease assay (**Supplementary Figure 5a**).

**Figure 7.**
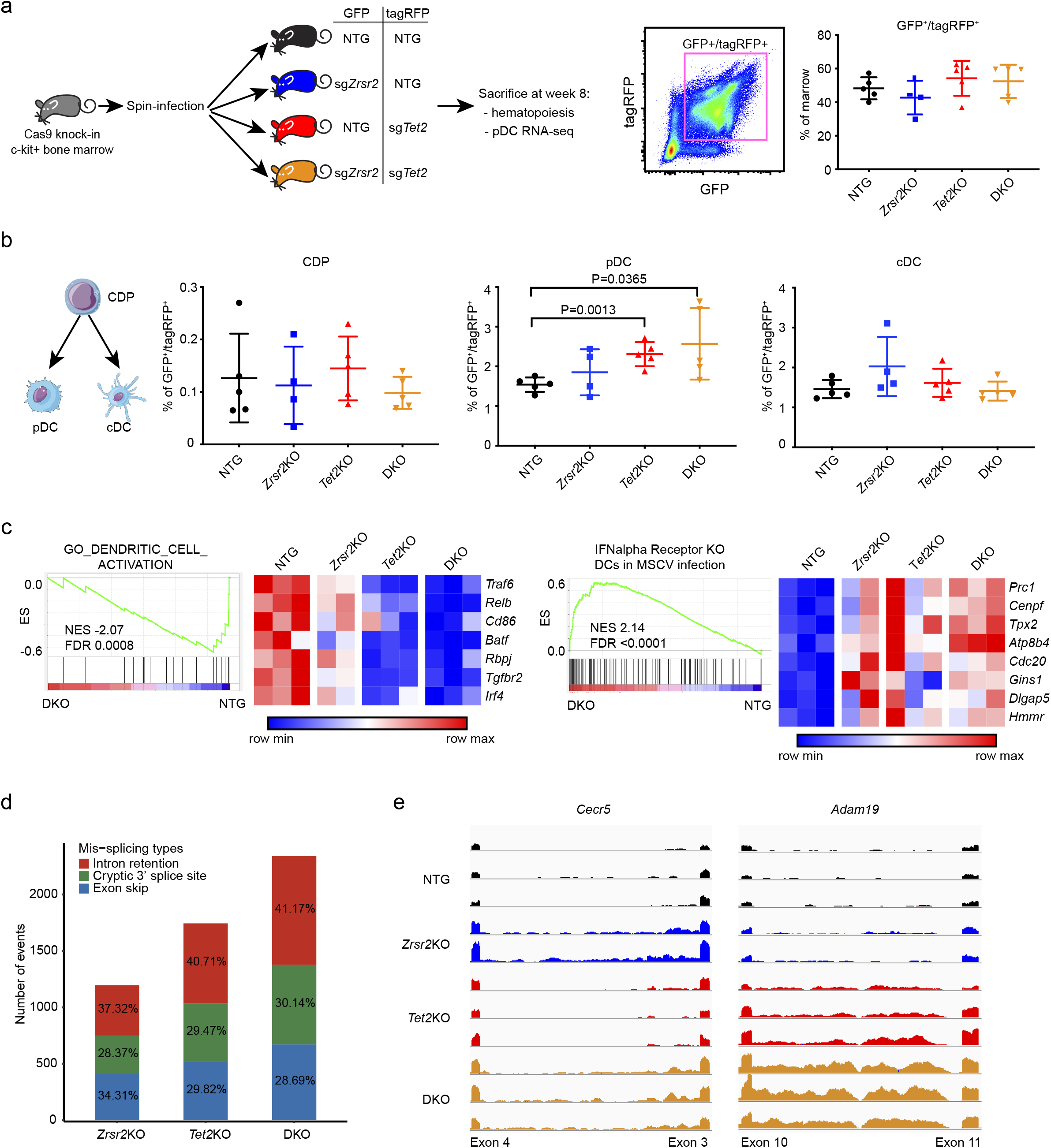
BPDCN-associated mutations in hematopoietic progenitors affect dendritic cell differentiation, RNA splicing, and activation signatures in vivo. **a.** Schematic of the in vivo experiment. sgRNA-transduced Cas9 knock-in bone marrow c-kit^+^progenitors were injected into lethally irradiated wild-type recipient mice. Each guide was marked with either GFP or tagRFP. The percentage of double positive cells in recipient bone marrow 8 weeks after transplantation is shown. **b.** Flow cytometry analysis of dendritic progenitor and mature subsets in sgRNA positive marrow cells, groups compared by t test. CDP, common dendritic progenitor; pDC, plasmacytoid dendritic cell; cDC, conventional dendritic cell. **c.** GSEA of RNA-seq in control (NTG) and *Zrsr2/Tet2-targeted* (DKO) pDCs showing changes in signatures related to DC activation and IFN alpha receptor-dependent gene expression in the setting of viral infection. Heatmap shows expression of the leading edge genes in GSEA from low (blue) to high (red). **d.** Mis-splicing events in *Zrsr2, Tet2*, and *Zrsr2/Tet2-targeted* (DKO) pDCs. Events in each condition were calculated by pairwise comparisons between control knockout (n=3) and *Zrsr2* (n=2), *Tet2* (n=3), and *Zrsr2/Tet2-targeted* (DKO, n=3) biologically independent replicates. The number of the events in each condition is shown in a single bar. The colors indicate different event types, intron retention (red), cryptic splice site (green), and exon skip (blue). e. Representative RNA-seq reads in *Zrsr2, Tet2*, and *Zrsr2/Tet2-targeted* (DKO) pDCs visualized on the same scale for *Cecr5* (intron retention associated with *Zrsr2* loss) and *Adam19* (intron retention in *Zrsr2* and *Te*f*2*-targeted pDCs with additive retention in *Zrsr2/Tet2-targeted* DKO pDCs).

We compared the GFP/tagRFP double positive populations in each of the four recipient groups (sgControl/sgControl, sg*Zrsr2*/sgControl, sgControl/sg*Tet2*, and *sgZrsr2/sgTet2)*. We focused on dendritic cells and their precursors, but we also analyzed other hematopoietic stem and progenitor (HSPC) populations. *Zrsr2* and *Tet2* targeting was associated with relative expansion of lineagenegative, Sca1+, cKit+ (LSK) HSPCs, common myeloid progenitors (CMP), and megakaryocyte-erythroid progenitors (MEPs) in the bone marrow (**Supplementary Figures 5b-c, 6**). In dendritic cell development, common dendritic progenitors (CDP) differentiate into conventional DCs (cDCs) and pDCs. In recipients of transduced and transplanted bone marrow, CDP and cDC percentage and absolute number were not different in *Zrsr2, Tet2*, or *Zrsr2/Tet2* double mutants compared to controls. In contrast, pDCs were modestly expanded, particularly in *Tet2* and *Zrsr2/Tet2* double mutant bone marrow (**Figure 7b**).

We assessed the transcriptome of mutant cells by sorting pDCs from bone marrow and performing RNA-seq. GSEA in mutant pDCs was negatively correlated with gene sets associated with DC activation and positively correlated with signatures of interferon alpha receptor knockout DCs exposed to viral infection (**Figure 7c**). This suggests that mutant mouse pDCs, similar to *ZRSR2*-mutant BPDCN cells, harbored signatures of decreased pDC activation and impaired type 1 interferon-dependent signaling. We also detected a higher frequency of mis-splicing events in the presence of *Zrsr2, Tet2*, and *Zrsr2/Tet2* mutation. Similar to what we observed in BPDCN, pDCs harboring these mutations accumulated several types of aberrant splicing (**Figure 7d**). Comparison of each group revealed genotype-specific patterns, such as *Zrsr2* mutation associated with a bias toward U12-type intron retention events, as we observed in human BPDCN cells with *ZRSR2* mutations (**Supplementary Figure 5d**). Mutation of *Tet2* alone was associated with splicing abnormalities and, in some genes, we observed additivity between *Tet2* and *Zrsr2* mutation to promote increased intron retention (**Figure 7e**). These data are consistent with existing evidence that mutations in *TET2* and other epigenetic modifiers can themselves promote splicing abnormalities and cooperate with splicing factor mutations to produce cooperative effects on splicing and hematopoiesis [31, 43]. Together, these in vivo data supported our findings in human BPDCN cells that ZRSR2 loss promotes specific transcriptome changes that provide a clonal advantage to pDCs with impaired activation.

## DISCUSSION

Our findings contribute to understanding how acquired mutations in hematopoietic cells drive dendritic cell transformation. By sorting tumor cells, we are confident that the somatic mutations reported are not simply passengers of concomitant myeloid malignancies, but indeed are present in the BPDCN cells. Furthermore, the findings presented here provide evidence that these mutations, particularly in RNA splicing factors and *TET2*, may have cell-type specific effects on pDCs, impairing their activation and apoptosis in response to TLR stimulation. We propose that innate immune signaling via IRF7 functions as a tumor suppressor in the pDC lineage to prevent malignant transformation. Further work will be required to determine if and how other mutations in BPDCN target this pathway and contribute to similar phenotypes.

The high frequency of *ZRSR2* loss-of-function mutations in BPDCN exclusively in male patients, nominate it as a lineage-specific Escape from X Inactivation Tumor Suppressor (EXITS) gene. By conservative estimation including only unambiguously inactivating alterations, *ZRSR2* mutations are associated with nearly half the male bias of BPDCN. We did not identify *ZRSR2* as an EXITS gene in our prior study [24] because it is most often mutated in BPDCN, MDS, and other myeloid neoplasms such as CMML, cancers that were not included in large-scale sequencing efforts like The Cancer Genome Atlas (TCGA). Therefore, this suggests that additional sex-biased cancer genes remain discoverable in understudied diseases. These data also illustrate how identifying functional consequences of a sex-biased mutation may uncover more generalized tumor biology that is relevant even in cases without the specific mutation of interest.

The effect of ZRSR2 loss was much more dramatic in TLR-stimulated cells than at baseline. The fact that *IRF7* is regulated by delayed splicing during inflammation [40] may help explain how mutations in multiple splicing factors can contribute to a similar phenotype in BPDCN. We previously found that *IRF7* was among the key pDC genes downregulated in BPDCN using single cell RNA-seq, suggesting that suppression of this pathway might be a common feature of pDC transformation even in the absence of a splicing factor mutation [14]. We would be surprised if all the activation defects observed in BPDCN are solely explainable by *IRF7* splicing abnormalities. However, it is plausible that distinct acquired alterations in pDCs converge on TLR signaling more generally to confer a clonal advantage by evasion of activation-induced apoptosis. Thus, optimal modeling of transformation by BPDCN-associated mutations may need to be performed in the setting of inflammation and/or in vivo.

*TET2, ASXL1*, and RNA splicing factor mutations are associated with clonal hematopoiesis (CH) and myeloid malignancies, such as MDS and CMML, that can pre-date BPDCN [44-46]. The pathogenesis model proposed here suggests that specific mutations in CH/MDS/CMML contribute to a pDC pool that is poised for transformation. A testable hypothesis for future research is that patients with MDS who develop BPDCN are enriched for having a history of abnormal inflammation. These findings also suggest that mutations in patients with myeloid malignancies or in individuals with age-related clonal hematopoiesis (ARCH or clonal hematopoiesis of indeterminate potential, CHIP) may affect pDC response to TLR stimulation and thereby contribute to impaired viral or tumor immunity. In support of this hypothesis, TET2 regulates *Irf7* expression via DNA methylation and is important for pDC type 1 interferon production and survival during viral infection in a mouse model [47]. Similarly, defective type 1 interferon production by pDCs is linked to deleterious *IRF7* variants in patients with inferior outcomes during severe COVID-19 / SARS-CoV-2 infection, a subgroup that is also male-biased and older than the general population [48]. Our data suggest that age-associated clonal hematopoiesis mutations, such as in *TET2* or *ZRSR2*, might also contribute to these hypoactive pDC phenotypes.

Finally, there may be therapeutic implications of these data. Splicing modulator drugs are in development and malignancies with splicing factor mutations are more sensitive to these agents [49]. Patients with BPDCN, stratified by presence of a splicing factor mutation or RNA mis-splicing pattern, could be included in splicing modulator clinical trials. Also, the UV mutational signature enrichment we observed may indicate that BPDCN would be a candidate for immune checkpoint blockade (ICB). ICB is only modestly active in other myeloid malignancies [50] but the cancers tested (AML, MDS) do not have a UV signature. In contrast, UV signatures predict response to checkpoint blockade in solid tumors, possibly due to qualitative differences in tumor neoantigens [51]. Furthermore, PD-L1 protein is expressed in approximately half of BPDCNs [52], which also supports clinical evaluation of ICB via PD-1/PD-L1 in the disease. Finally, we previously found that BPDCN is highly dependent on BCL2 and sensitive to the BCL2 inhibitor venetoclax [32]. We do not yet know if this is related to splicing mutations and impaired activation-induced apoptosis, but the data presented here provide additional rationale for evaluating BCL2 inhibition in BPDCN.

Genes mutated in BPDCN are similar to those in MDS and AML. However, many clinical characteristics of BPDCN and AML are distinct, including epidemiology, clinical presentation, pathology, and response to certain therapies. These data suggest a mechanism by which the consequences of the same gene mutation (the ‘seed’; e.g., *ZRSR2)*, could have lineage-specific effects based on cell context (the dendritic cell ‘soil’). The TLR-IRF7-type 1 interferon-apoptosis axis that is dysfunctional in BPDCN may not be the target of the same splicing mutations in other cancers. This highlights the need to study BPDCN and other rare cancers as unique entities and to consider the importance of cellular milieu when evaluating the function of cancer genes.

## ACKNOWLEDGEMENTS

This work was supported by the Sumitomo Life Welfare and Culture Foundation (KT), the National Cancer Institute R37 CA225191 (AAL) and R35 CA231958 (DMW), the Damon Runyon Cancer Research Foundation (GKG), the Mark Foundation for Cancer Research (AAL), Doris Duke Charitable Foundation (AAL), Ludwig Center at Harvard (AAL), and the American Society of Hematology (AAL).

## DISCLOSURES

AAL has received research funding from AbbVie and Stemline Therapeutics and consulting fees from N-of-One and Qiagen. GKG has received research funding from Calico Life Sciences and consulting fees from Moderna Therapeutics. MS and SB were employees of H3 Biomedicine at the time of this study. MS is currently an employee of Remix Therapeutics (Cambridge, MA).

## AUTHOR CONTRIBUTIONS

KT, SSC, VM, CMK, LCH, JT, SSK, GKG, MG, KB, YYL, FA-D, SB conducted experiments and analyzed data; MS, SB, HY analyzed data; SBL, AL Jr., EAM, FJ, PPP, DMW, PSH, MK, NP, OAW, HPK provided critical reagents and analyzed data; KT and AAL designed the study, interpreted the data, and wrote the paper; all authors participated in editing the paper.

## METHODS

### Cell lines

293T packaging cells were obtained from ATCC. CAL1 cells were provided by T. Maeda (Nagasaki University, Nagasaki, Japan). Cell line identity was verified by short tandem repeat (STR) profiling in the DFCI Molecular Diagnostics Core and cells were verified to be mycoplasma-free by regular testing at least every 6 months. CAL1 cells were cultured in RPMI 1640 supplemented with 10% fetal bovine serum (FBS), 1 x penicillin/streptomycin (Gibco, 15140122), and 1% GlutaMAX (Gibco, 35050061). 293T packaging cells were cultured in DMEM supplemented with 10% FBS, 1 x penicillin/streptomycin. Proliferation and apoptosis were measured at the indicated times starting with a concentration of 2×10^5^ cells/ml. Proliferation was measured by CellTiter-Glo (Promega, G7572) and caspase activity was measured by Caspase-Glo 3/7(Promega, G8092).

### Lentiviral infections

293T cells were transfected with 10.8 μg psPAX2, 2.4 μg pVSV-G, and 10.8 μg of a lentiviral expression vector with Lipofectamine 2000 (Thermo Fisher, 11668500). Viral supernatant was harvested 48 and 72-hours after transfection and concentrated by ultracentrifugation at 23,000g for 2 hours at 4°C. 2×10^5^ cells were infected in the presence of 1.5 ml of viral supernatant and 4 μg/ml polybrene (Santa Cruz Biotechnology, SC-134220).

### Mouses

All animal experiments were performed with approval from the Dana-Farber Cancer Institute (DFCI) Animal Care and Use Committee. C57BL/6J and Rosa-Cas9 knock-in mice (B6J.129(Cg)-Gt (ROSA)26Sor^tm1.1(CAG-cas9*,-EGFP)Fezh^/J) were purchased from Jackson Laboratory (026179). Cas9 knock-in mice were sacrificed and bone marrow cells were harvested from both legs, iliac bones and spine for transplantation. CD117^+^ cells were sorted by using magnetic beads (Miltenyi Biotec #130-094-224) after red cell lysis. After culture in StemSpan™ SFEM with 50 ng/ml SCF (Gold Bio Technology, 1320-01) and 50ng/ml TPO (Gold Bio Technology, 1320-06) overnight, CD117/c-kit^+^ cells were infected with virus particles in the presence of 4 μg/ml polybrene. Transduced cells were injected into the tail vein of lethally irradiated (5.5 Gy x2 split doses) C57BL/6J mice with rescue marrow. Animals were sacrificed 8 weeks after the transplantation. Bone marrow and splenocytes were harvested for analysis.

### Plasmids for cDNA and shRNA expression and CRISPR/Cas9 gene targeting

shRNAs were subcloned into a Tet-on pLKO-puro (Addgene, #21915) via AgeI and EcoRI restriction sites. shRNAs were induced with 1 μg/ml doxycycline after selecting transduced cells in 1 μg/ml puromycin. sgRNAs were subcloned into a lentiviral expression vector that coexpresses GFP (pLKO5.sgRNA.EFS.GFP; Addgene, #57822) or tagRFP (pLKO5.sgRNA.EFS.tagRFP; Addgene, #57823) via the BsmBI restriction site. Cell lines stably expressing the Cas9 nuclease were generated by infection with the empty lentiCRISPRv2 lentivirus (Addgene, #52961, not containing an sgRNA guide) using standard methods (https://www.addgene.org/viral-vectors/lentivirus/lenti-guide/). Cells were selected in puromycin and FLAG-Cas9 expression was confirmed by Western blot. Cas9-expressing cell lines were infected at a density of 2×10^5^ cells in 1.5 mL media in the presence of 4 μg/mL polybrene (Santa Cruz Biotechnology, SC-134220). sgRNA resistant human ZRSR2 DNA fragment was synthesized by Twist Bioscience and cloned into pRRL.idTomato.

*sgRNA/shRNA sequences*:

hZRSR2_sgRNA1, TGACGTTTCCCGAGAAACCA
hZRSR2_sgRNA2, ACAGTTCCTAGACTTCTATG
hZRSR2_sgRNA3, CTTCCAGCCACAAAAAGTAC
hZRSR2 shRNA, CAACAGTTCCTAGACTTCTAT
Control shRNA, CTCAGTTCCAGTACGGCTCCA
mZrsr2_sgRNA1, ACGTCTCTCCTTCTTCCGGA
mZrsr2_sgRNA2, GCATGAAGAATGGTTACTGA
mTet2_sgRNA1, GAATACTATCCTAGTTCCGAC
mTet2_sgRNA2, GAACAAGCTCTACATCCCGT
NTG1, GACGGAGGCTAAGCGTCGCAA
NTG2, GCGCTTCCGCGGCCCGTTCAA

*sgRNA resistant ZRSR2 cDNA sequence*:

ATGGCTGCGCCCGAGAAGATGACGTTTCCCGAGAAACCATCCCACAAAAAGTACAGGGCCGCCCTGAAGAAGGAGAAACGAAAGAAACGTCGGCAGGAACTTGCTCGACTGAGAGACTCAGGACTCTCACAGAAGGAGGAAGAGGAGGACACTTTTATTGAAGAACAACAACTAGAAGAAGAGAAGCTATTGGAAAGAGAGAGGCAAAGATTACATGAGGAGTGGTTGCTAAGAGAGCAGAAGGCACAAGAAGAATTCAGAATAAAGAAGGAAAAGGAAGAGGCGGCTAAAAAACGGCAAGAAGAACAAGAGAGAAAGTTAAAGGAACAATGGGAAGAACAGCAGAGGAAAGAGAGAGAAGAGGAGGAGCAGAAACGACAGGAGAAGAAAGAAAAAGAGGAAGCTTTGCAGAAGATGCTGGATCAGGCTGAAAATGAGTTGGAAAATGGTACCACATGGCAAAACCCAGAACCACCCGTGGATTTCAGAGTAATGGAGAAGGATCGAGCTAATTGTCCCTTCTACAGTAAAACAGGAGCTTGCAGATTTGGAGATAGATGTTCACGTAAACATAATTTCCCAACATCCAGTCCTACCCTTCTTATTAAGAGCATGTTTACGACGTTTGGAATGGAGCAGTGCAGGAGGGATGACTATGACCCTGACGCAAGCCTGGAGTACAGCGAGGAAGAAACCTACCAACAGTTCCTAGACTTCTATGAAGATGTGTTGCCCGAGTTCAAGAACGTGGGGAAAGTGATTCAGTTCAAGGTCAGCTGCAATTTGGAACCTCACCTGAGGGGCAATGTATATGTTCAGTACCAGTCGGAAGAAGAATGCCAAGCAGCCCTTTCTCTGTTTAACGGACGATGGTATGCAGGACGACAGCTGCAGTGTGAATTCTGCCCCGTGACCCGGTGGAAAATGGCGATTTGTGGTTTATTTGAAATACAACAATGTCCAAGAGGAAAGCACTGCAACTTTCTTCATGTGTTCAGAAATCCCAACAATGAATTCTGGGAAGCTAATAGAGACATCTACTTGTCTCCAGATCGGACTGGCTCCTCCTTTGGGAAGAACTCCGAAAGGAGGGAGAGGATGGGCCACCACGACGACTACTACAGCAGGCTGCGGGGAAGGAGAAACCCTAGTCCAGACCACTCCTACAAAAGAAATGGGGAATCCGAGAGGAAAAGTAGTCGTCACAGGGGGAAGAAATCTCACAAACGCACATCAAAGAGTCGGGAGAGGCACAATTCACGAAGCAGAGGAAGAAATAGGGACCGCAGCAGGGACCGCAGCCGGGGCCGGGGCAGCCGGAGCCGGAGCCGGAGCCGGAGCCGCAGGAGCCGCCGCAGCCGGAGCCAAAGTTCCTCTAGGTCCCGAAGTCGTGGCAGGAGGAGGTCGGGTAATAGAGACAGAACTGTTCAGAGTCCCAAATCCA AATAA

### Flow cytometry

Cells were washed with PBS containing 2% FBS before staining and were then incubated with the indicated antibodies indicated for 30 minutes in the dark at 4°C followed by a final wash in PBS containing 2% FBS and analyzed. In the case of murine samples, we lysed red blood cells prior to staining. Antibodies for staining are indicated in the following table. Gating strategy for bone marrow cells are defined as indicated below and in **Supplementary Figure 6**. Flow cytometry was performed using a BD LSRFortessa™ X-20 and analyzed using FlowJo software, version 10.

#### Antibodies for flow cytometry

|  | Target | Fluorophore | Provider, Catalog number | Dilution |
| --- | --- | --- | --- | --- |
| Mouse | CD11b | APC/Cy7 | Biolegend, #101226 | 1/200 |
|  | Gr1 | PerCP/Cy5.5 | BioLegend, #108427 | 1/100 |
|  | CD19 | Brilliant Violet 605 | BioLegend, #115540 | 1/100 |
|  | CD3 | Brilliant Violet 421 | BioLegend, #100228 | 1/100 |
|  | CD11c | Biotin | BioLegend, #117303 | 1/100 |
|  | MHC class 2 | Alexa700 | BioLegend, #107621 | 1/100 |
|  | B220 | APC/Cy7 | BioLegend, #103224 | 1/100 |
|  | CD317 | Brilliant Violet 421 | BioLegend, #127023 | 1/100 |
|  | SiglecH | PerCP/Cy5.5 | BioLegend, #129613 | 1/100 |
|  | Lineage Cell Detection Cocktail | Biotin | Miltenyi Biotec, #130-092-613 | 1/100 |
|  | CD117 | Brilliant Violet 711 | BioLegend, #105835 | 1/100 |
|  | CD115 | PE/Cy7 | BioLegend, #135523 | 1/100 |
|  | CD135 | PE/Cy5 | BioLegend, #135311 | 1/100 |
|  | CD127 | APC/Cy7 | BioLegend, #135039 | 1/100 |
|  | Sca1 | Brilliant Violet 605 | Fisher Scientific, #BDB744324 | 1/100 |
|  | CD34 | Alexa700 | Fisher Scientific, #BDB560518 | 1/100 |
|  | CD16/32 | PerCP/Cy5.5 | BioLegend, #101323 | 1/100 |
|  | Streptavidin | Brilliant Violet 510 | BioLegend, #405233 | 1/100 |
|  | Streptavidin | PE/Cy7 | BioLegend, #405206 | 1/100 |
| **Human** | CD80 | PE | Fisher Scientific, #BDB557227 | 1/100 |
|  | CD123 | APC | BD Biosciences, #658172 | 1/100 |
|  | BDCA2 | Brilliant Violet 785 | BioLegend, #354221 | 1/100 |
|  | CD3 | FITC | BioLegend, #300406 | 1/100 |
|  | CD45 | Pacific Blue | BioLegend, #103230 | 1/100 |
|  | CD4 | PE | BD Biosciences, #555347 | 1/100 |
|  | CD56 | PE/Cy7 | BD Biosciences, #557747 | 1/100 |

#### HSPC markers by population of interest

| **Population** | **Surface markers for flow cytometry** |
| --- | --- |
| LSK | Lin-, CD117+, Sca-1+ |
| LK | Lin-, CD117+, Sca-1- |
| CMP (common myeloid progenitor) | Lin-, CD117+, Sca-1-, CD34+, CD16/32- |
| GMP (granulocyte-monocyte progenitor) | Lin-, CD117+, Sca-1-, CD34+, CD16/32+ |
| MEP (megakaryocyte-erythroid progenitor) | Lin-, CD117+, Sca-1-, CD34-, CD16/32- |
| CDP (common dendritic progenitor) | Lin-, CD127-, CD115+, CD135+, CD117-dim |
| CLP (common lymphoid progenitor) | Lin-, CD135+, CD117-dim, CD127+ |
| MDP (monocyte-dendritic progenitor) | Lin-, CD127-, CD115+, CD135+, CD117+ |
| pDC | CD19<sup>-</sup>, B220<sup>+</sup>, CD317<sup>+</sup>, SiglecH<sup>+</sup> |
| cDC | MHC class II<sup>+</sup>, CD11b<sup>+</sup>, CD11c<sup>+</sup> |

### Western blotting

Samples were prepared by lysing in RIPA buffer (Boston BioProducts, BP-115) with protease inhibitor cocktail (Thermo Fisher Scientific, 862209) and sonicated before quantification by BCA assay (Thermo Fisher Scientific, 23225). Samples were prepared with SDS sample buffer (Boston BioProducts, BP-110R), boiled for 10 minutes at 98°C, and recovered by spinning at 12,000*g* for 5 minutes at 4°C before loading onto the gel. The gel was run for 80 minutes at 120V in SDS running buffer (Boston BioProducts, BP-177) before being transferred to a PVDF membrane (Thermo Fisher Scientific, 7462) for 7 minutes at 20 V using iBlot 2 (Thermo Fisher Scientific). Blots were blocked in 5% dried milk (AppliChem, A0830) in tris-buffered saline (TBS) with 0.1% Tween-20 for 1 hour before being incubated overnight in antibodies recognizing ZRSR2 (kindly provided by Dr. Michael Green, University of Massachusetts Medical School, Massachusetts, 1:2000), IRF7 (Cell Signaling technology, #4920, 1:1000), TRAIL (Cell Signaling technology, #3219, 1:1000), β-actin (Sigma-Aldrich, A5441, 1:10,000). The blots were washed 3 times in TBS with 0.1% Tween-20 before being incubated with either rabbit (Santa Cruz Biotechnology, SC-2004) or mouse (Santa Cruz Biotechnology, SC-2005) secondary HRP-conjugated antibodies before being imaged using ECL substrate (Bio-Rad, 170-5061) on an ImageQuant LAS-4000 (GE Healthcare, 28-9607-59AB).

### T7 endonuclease (T7E1) assay

Genomic DNA from mouse bone marrow cells was extracted by the DNeasy Blood and Tissue Kit (Qiagen, #69504). The DNA region containing sgRNA target sites was amplified by PCR using specific primers (*Zrsr2* Forward, CCCATGGCATCTTTGTCTATAATCT-; *Zrsr2* Reverse, GCTCAGCTAAGCACTTACTCAATG; *Tet2*_Forward, ACATACTCCTCAGACGCAGG; *Tet2*_Reverse, CTGGCATGTACCTGGATTGC). PCR products were purified with NucleoSpin Gel and PCR Clean-Up (Takara Bio, #740609). 400 ng of PCR products mixed with NEB buffer 2 were denatured at 95°C for 5 minutes and ramped down to 25°C. T7 Endonuclease 1 (NEB, M0302) was added to the mixture and incubated at 37°C for 15 minutes. 0.25 M EDTA was added to stop the reaction and samples were analyzed on 2% agarose gel.

### RT-PCR

RNA was extracted from cell lines using TRIzol (Life Technologies, 15596018) and quantified via NanoDrop. The High-Capacity cDNA Reverse Transcriptase Kit (Thermo Fisher Scientific, 4368814) was used to generate cDNA. qRT-PCR was performed for human *IRF7*, using *GAPDH* as the internal control, using the primers as below, and SYBR Green PCR master mix per the manufacturer’s instructions (Thermo Fisher Scientific, 4367659). Relative quantification was calculated using the ΔΔCt method. *IRF7* For, TACCATCTACCTGGGCTTCG; *IRF7* Rev, GAAGACACACCCTCACGCTG; *GAPDH* For, GCACCGTCAAGGCTGAGAAC; *GAPDH* Rev, TGGTGAAGACGCCAGTGGA. For RT-PCR, the following primers were used: *DERL3* For, GGCCGACTTCGTCTTCATGTTTC; *DERL3* Rev, CAGGTCCACGAGGATGGAGT.

### Cytokine quantitation

Peripheral blood mononuclear cells (PBMCs) were isolated by density gradient centrifugation (Ficoll®-Paque PLUS, GE Healthcare, 17-1440-03) from healthy donors. Normal pDCs were enriched by using magnetic beads (Miltenyi Biotec, 130-097-415). CD123^+^BDCA2^+^ cells were sorted for normal pDC and CD45^dim^CD123^+^ cells were sorted for BPDCN PDX tumor cells using a FACSAria. CAL1 cells were seeded at 2×10^5^/ml, BPDCN tumor cells and normal pDCs were seeded at 1×10^5^/ml. Cells were stimulated with either LPS (SigmaAldrich, L3129) or R848 (Invitrogen, tlrl-r848) for 24 hours. Supernatants were collected and cytokines were measured by either Bio-Plex Pro (BIO RAD, #17AL001M) or ProcartaPlex (Thermo Fisher, PPX-24).

### Whole exome sequencing and analysis

All patients were consented to a local institutional review board (IRB) approved protocol. Tumor and paired germline DNA samples were collected and prepared for whole-exome sequencing using the Agilent SureSelect Human All Exon V5(50M) capture. Sequencing was performed on Hiseq4000 platform with a 150 base pair paired-end protocol targeted to 100x mean coverage. The sequences were generated as 90/100 base paired-end reads using Illumina base calling Software (ver. 1.7). The adapter sequences and low-quality reads were filtered from the raw sequencing data and the “clean data” was aligned by Burrows-Wheeler Aligner (BWA) [53] with the reference of human genome build37 (hg19). The BAM files were validated by steps of fixing mate information of the alignment, adding read group information and removing duplicate reads, and were then applied to variant calling analysis using GATK toolkit [54]. The reference (hg19) and germline variant sites were used to detect somatic or tumor-specific mutations and copy number alterations. The details of the methods in GATK toolkit are described in [55, 56]. The identified BPDCN somatic mutations were analyzed with those reported for acute myeloid leukemia (AML) and skin melanoma [57] for presence of previously defined global DNA mutational signatures in the Catalog of Somatic Mutations in Cancer (COSMIC) database [20] using the R package, “MutationalPatterns” [58].

### RNA sequencing and analysis

Samples from BPDCN patients, PDX, CAL1 and mouse models were collected for total RNA sequencing using the Arcturus PicoPure RNA Isolation Kit (Life Technologies). Libraries were prepared using the Ovation kit (Nugen, #0340) using 50 ng of input total RNA and 20 cycles of amplification. The Illumina Hi-Seq platform was used to generate single or paired-end sequencing results. The raw FASTQ data were analyzed by VIPER pipeline [59] which combines STAR [60] alignment (mapped to hg19 or mm10) and gene expression analysis such as unsupervised clustering, PCA based on Cufflinks [61] and differential expression with DESeq2 [62]. The expression data were used for pathway enrichment analysis with GSEA [63, 64].

### Differential splicing analysis

Splicing events were identified as described in [25]. The method calculates a mis-splicing index (MSI) for each splicing event in comparison to the reference genome (human hg19 or mouse mm10) and classifies them as intron retention, exon skip, and incorrect splice site usage. The difference in MSI between two samples was used for direct comparisons by defining the delta MSI (ΔMSI) as the difference in MSI at a given site between two samples. The statistical significance of differences in splicing events was evaluated by Fishers’ exact test with adjusted P-value for multiple hypothesis testing. Differences with ΔMSI>0.2 and P≤0.05 were considered significant.

### Data visualization and availability

Analysis results were visualized using R 3.5.2 (R Foundation for Statistical Computing) [65] and Python (https://www.python.org). Co-mutation plot was created using the “GenVisR” package in R [66]. RNA sequencing data will be deposited in GEO, accession number pending.

---

**Supplementary Figure 1.**
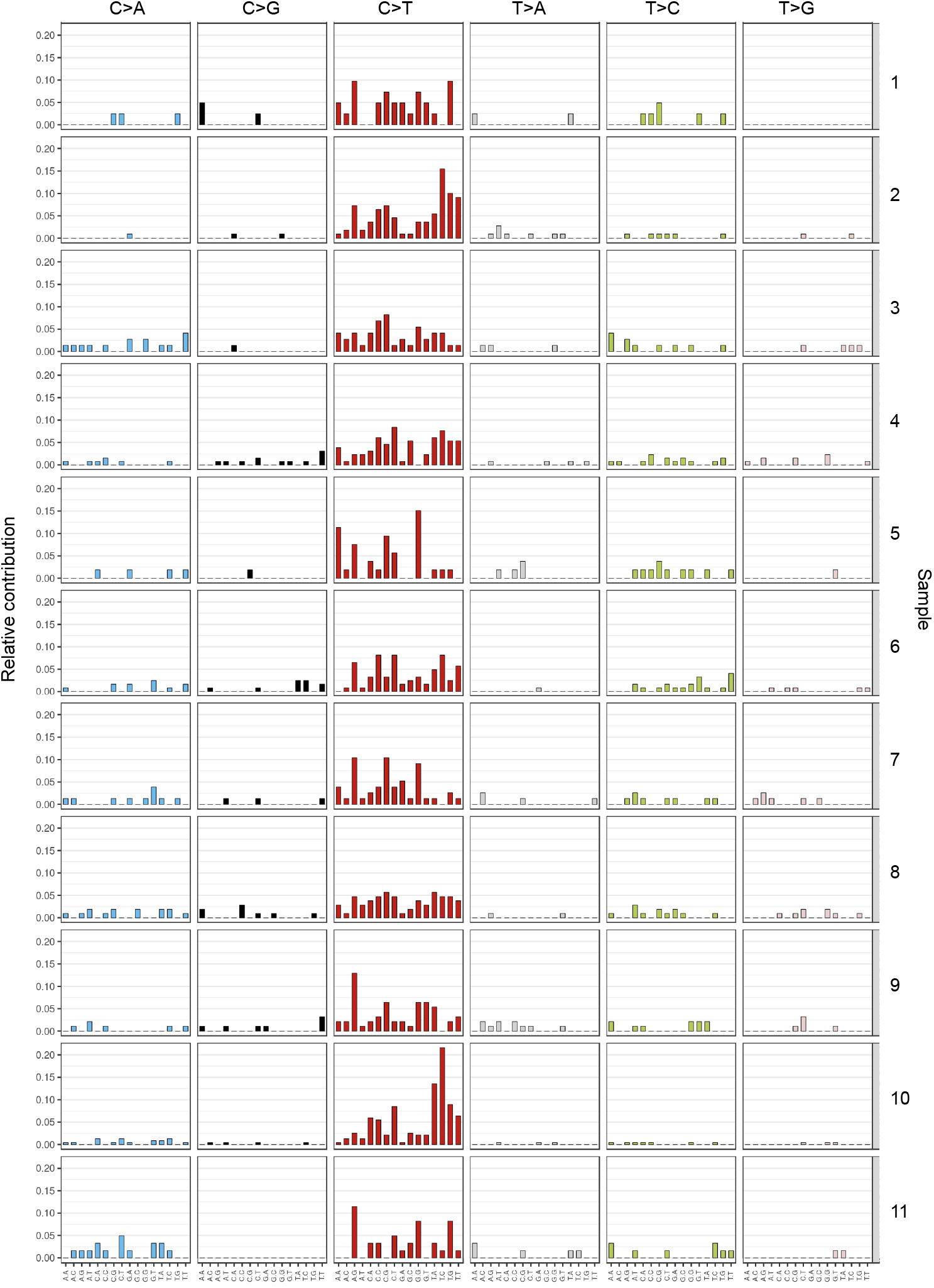
Global mutational patterns in BPDCN. Somatic DNA mutations in 11 BPDCN whole exomes compared to matching germline are shown as patients by row and base change in columns plotted as relative contribution to the total number of base changes. Mutations are grouped by the specific base change on either strand (C>A, C>G, C>T, T>A, T>C, or T>G) and each individual bar plot represents the bases flanking the mutated base. Signatures of UV-induced mutational processes are characterized by a preponderance of C>T substitutions at dipyrimidine sites, such as CC>TT.

**Supplementary Figure 2.**
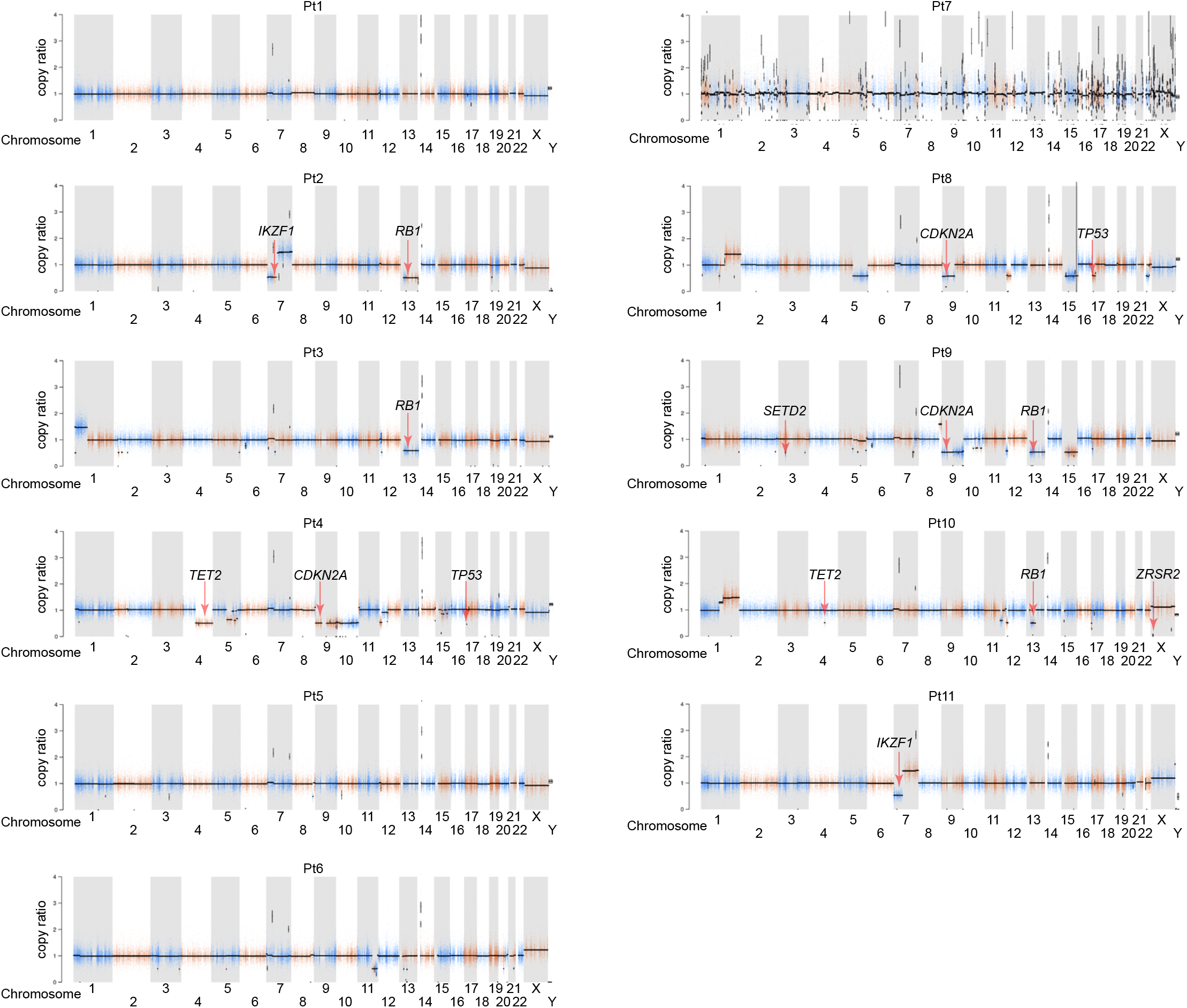
Somatic alterations in DNA copy number from isolated BPDCN cells. DNA copy ratio of tumor to normal calculated from whole exome sequencing is plotted for 11 BPDCN tumor/normal pairs by chromosome, where a copy ratio of 1 means no difference between tumor and normal, <1 is copy loss and >1 is copy gain. Chromosome positions of genes of interest are indicated.

**Supplementary Figure 3.**
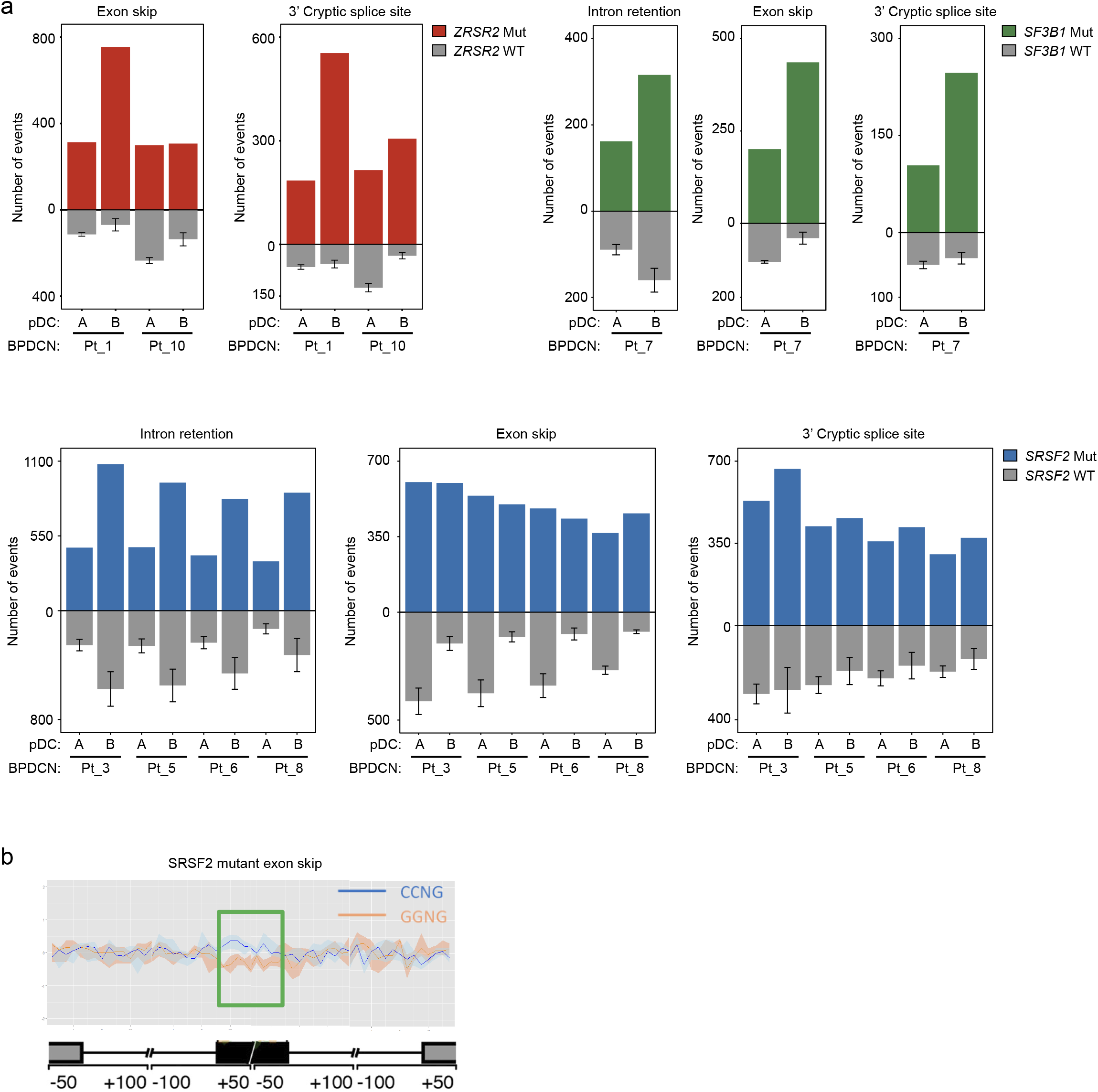
Aberrant RNA splicing events in BPDCN by genotype compared to pDCs. **a.** Mis-splicing events in the indicated BPDCN samples compared to normal donor pDCs are shown, categorized as intron retention, exon skipping, or cryptic 3’ splice site events. BPDCNs with *ZRSR2* mutation (n=2), BPDCN with an *SF3B1* mutation, BPDCNs with *SRSF2* mutation (n=4), and BPDCN patients without any known splicing mutation (n=4) are shown, using P≤0.05 in the respective splicing factor mutant cases compared to normal pDCs (two independent samples, A and B) to define events. **b.** Meta-gene tracing for an *SRSF2* mutated BPDCN showing the canonical exon inclusion/exclusion pattern averaged across the genome for an exon containing a CCNG (inclusion favored) or GGNG (exclusion favored) SRSF2 RNA recognition motif (RRM), similar to that previously defined for *SRSF2* mutations in MDS [30].

**Supplementary Figure 4.**
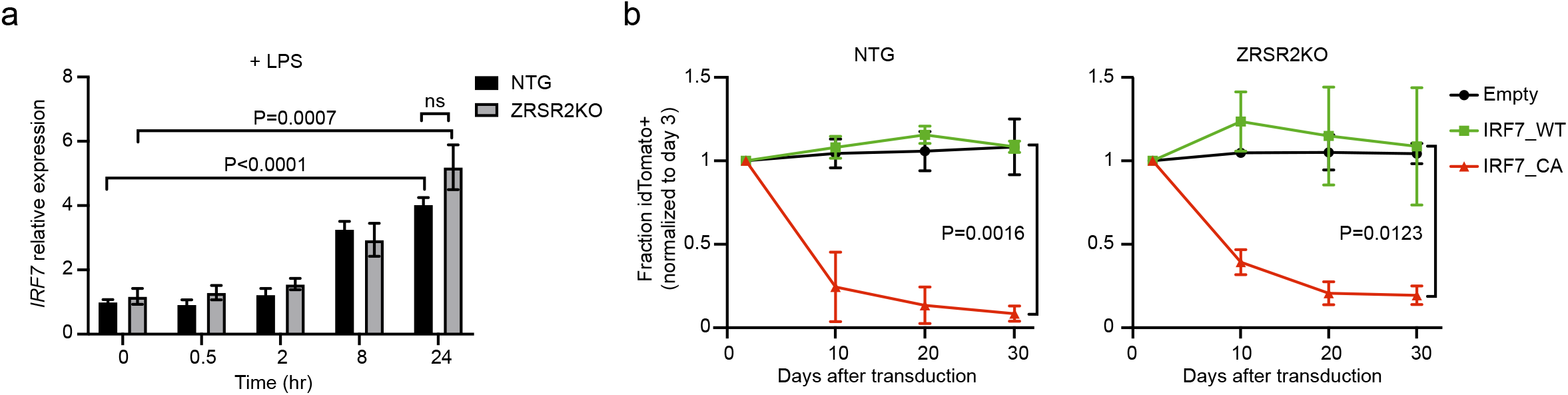
IRF7 RNA induction after LPS stimulation and growth disadvantage after constitutive IRF7 activation are similar in control or *ZRSR2* knockout cells. **a.** Induction of *IRF7* RNA relative to *GAPDH* in both control and *ZRSR2* knockout CAL1 cells is shown at the indicated times after stimulation with LPS. Each group represents biological triplicates compared by t test. **b.** idTomato positive cells relative to day 3 after transduction with empty vector, wild-type IRF7 (IRF7_WT), or constitutively active IRF7 (IRF7_CA) in CAL1 cells with control sgRNA or knockout of *ZRSR2*. Each group represents biological triplicates with IRF_WT and IRF_CA compared by t test.

**Supplementary Figure 5.**
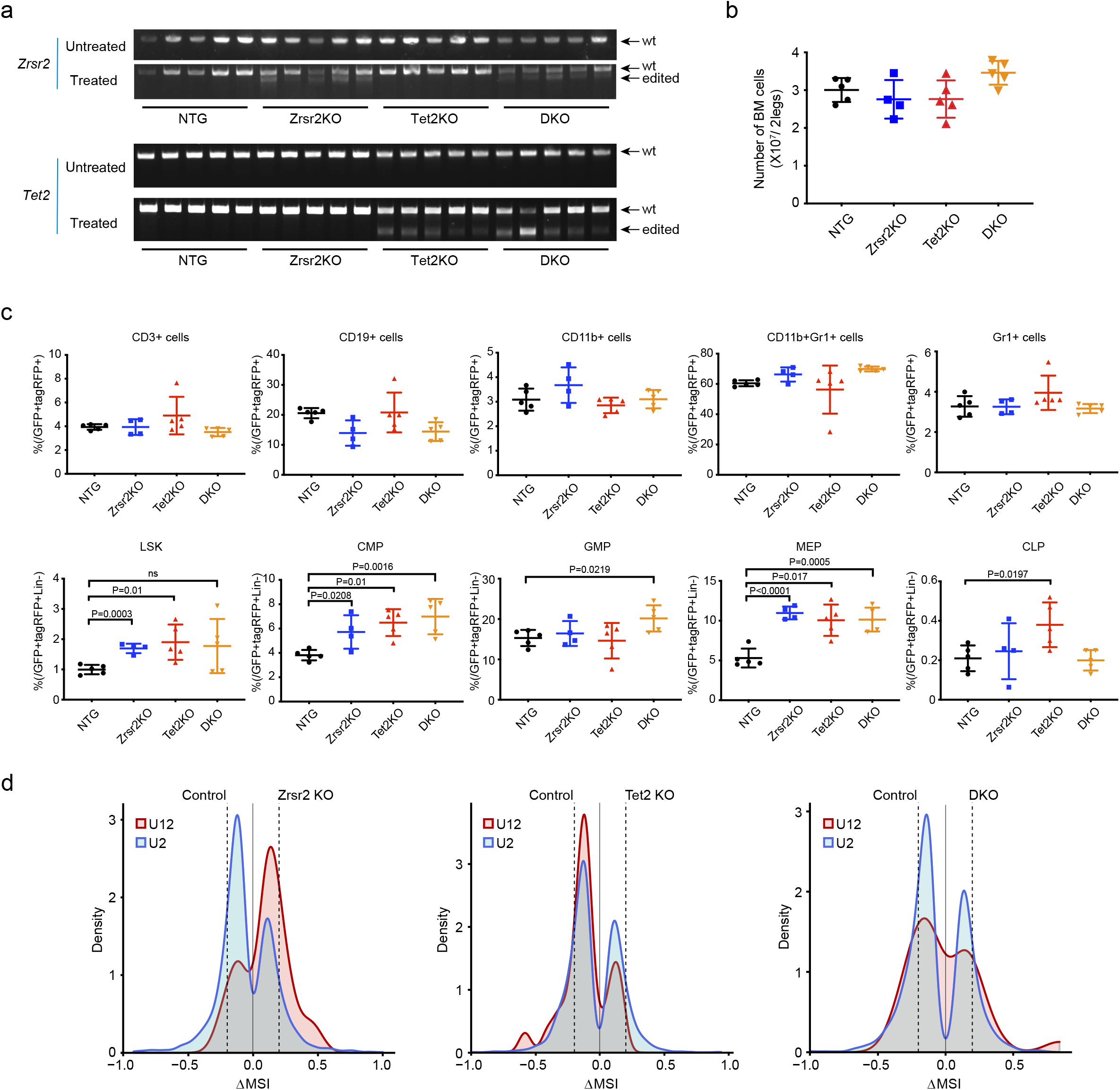
In vivo consequences of *Zrsr2* and *Tet2* CRISPR/Cas9 targeting on hematopoiesis and pDC intron retention. **a.** T7 endonuclease assay on bone marrow cells from transplant recipients to detect indel mutations after CRISPR/Cas9 editing. Amplified sgRNA target sites in *Zrsr2* or *Tet2* from cells harvested from the indicated groups (n=5 independent animals/genotype) are indicated. “Untreated” and “Treated” refer to whether the PCR products were incubated with T7 endonuclease. **b.** Total number of nucleated cells per two legs (femurs and tibias) in the bone marrow of recipient mice of the indicated CRISPR/Cas9 edited transplants. **c.** Bone marrow hematopoietic populations as percentage of GFP/tagRFP double positive cells in the bone marrow (top row) or of Lineage-negative GFP/tagRFP double positive cells (bottom row). Groups compared by t test. LSK, lineage-Sca1+cKit+; CMP, common myeloid progenitor; GMP, granulocyte-monocyte progenitor; MEP, megakaryocyte-erythroid progenitor; CLP, common lymphoid progenitor. **d.** Density plots of frequencies of U2-type (blue) and U12-type (red) introns among aberrantly retained introns (Fisher’s exact test p-value ≤ 0.05) in pairwise analyses of GFP+/tagRFP+ pDCs from the indicated genotypes versus non-targeting control sgRNA-containing pDCs.

**Supplementary Figure 6.**
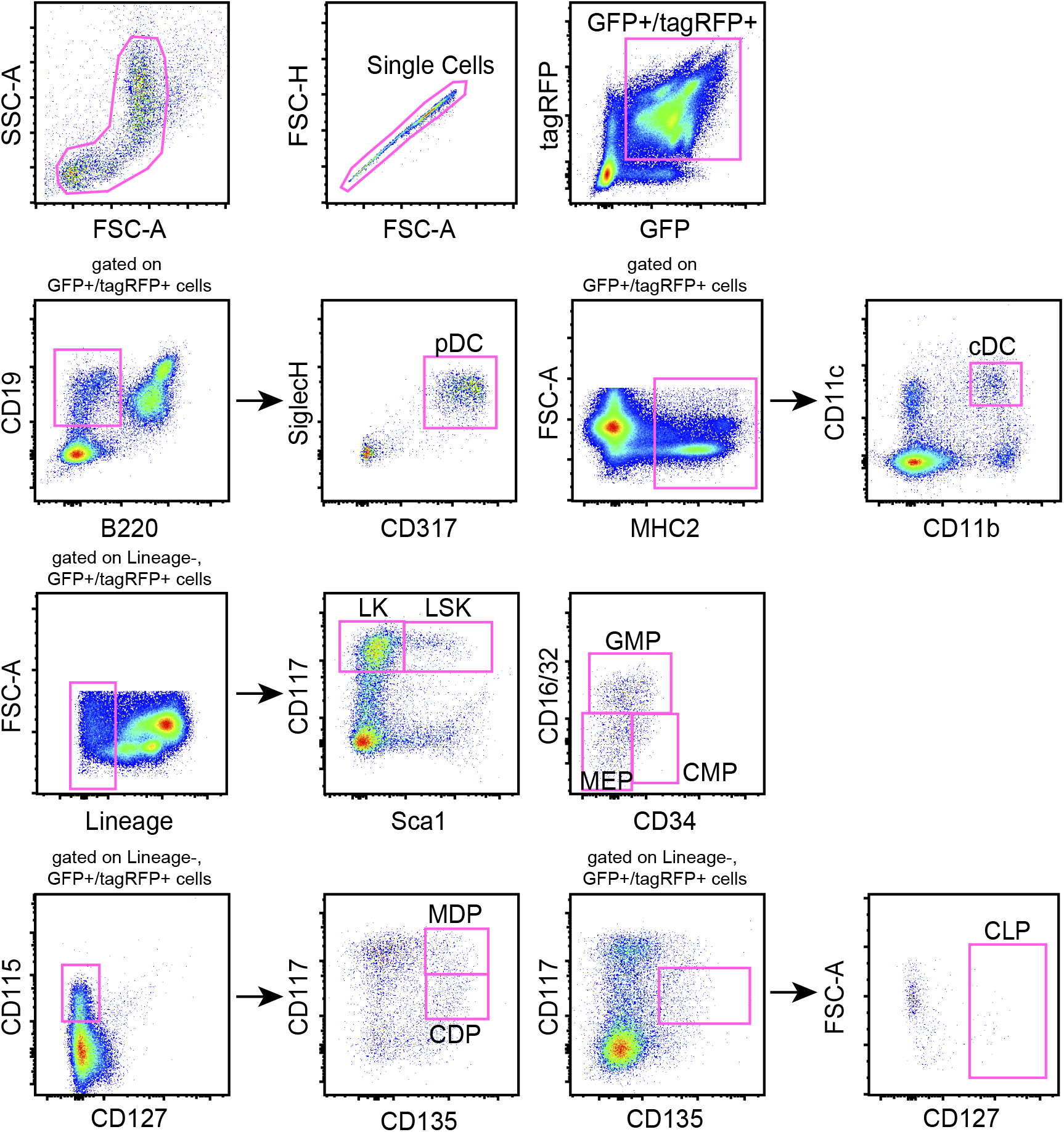
Gating strategy for bone marrow HSPC and DC subpopulations. Representative flow cytometry plots are shown for mouse bone marrow to define the gating strategy used to quantify hematopoietic stem and progenitor cells (HSPCs) and mature hematopoietic cells, including cDCs and pDCs.

